# Therapeutic vulnerabilities of the drug-resistant fungal meningitis pathogen *Cryptococcus neoformans*

**DOI:** 10.64898/2026.10.31.763343

**Authors:** Manning Y. Huang, Patrick Hecht, Kyle Cesar, Massimiliano Ciranni, Angela L. Wei, Michael J. Boucher, Sydney L. Briner, Léa Lortal, Veda Nayak, Ankur Ankur, Tuo Wang, Tamara L. Doering, Vito Paolo Pastore, Hiten D. Madhani

## Abstract

Treatment options are limited for the priority fungal pathogen *Cryptococcus neoformans*. We describe CRISPR-insert, a scalable gene inactivation method for global interrogation of drug targets. Genome-wide screens using this methodology combined with machine learning predict 1,297 essential genes under liberal criteria. Unexpectedly absent from this list is *FKS1*, encoding the target of echinocandin drugs to which *C. neoformans* is resistant. Contrary to longstanding literature, targeted deletion shows that *FKS1* is not essential. Solid-state NMR reveals that *fks1*Δ mutants lack β-1,3-glucan, arguing against compensation by another synthase. We experimentally test essentiality predictions for genes lacking human orthologs, identifying high-priority drug target candidates. To uncover targets for combination therapies, we establish genome-scale quantitative genetic interaction mapping. We use this method to uncover synthetic lethal dependencies of echinocandin resistance determinants, including the druggable calcineurin and Cdc50-Apt1 pathways. CRISPR-insert illuminates therapeutic vulnerabilities of an invasive fungal pathogen.

## INTRODUCTION

It is estimated that 6.5 million invasive fungal infections occur worldwide each year, with 3.8 million associated deaths,^1^ and invasive fungal infections in the United States cost roughly $19.4 billion annually^2^. Most human fungal pathogens are opportunistic and cause severe outcomes in immunocompromised patients. Because fungi are closely related to humans, fewer antifungal drugs exist than antibacterials, and antifungals can be highly toxic. For example, the side effects of intravenous amphotericin B, an antifungal used as first-line therapy for severe infections, include potent nephrotoxicity and seizures^3^.

*Cryptococcus neoformans* is a top-ranked WHO priority fungal pathogen, estimated to cause 194,000 infections with 147,000 associated deaths every year and to account for nearly 20% of AIDS-related deaths^1,4^, due to meningoencephalitis that is invariably fatal without treatment^5^. Other immunocompromised patient populations are also highly susceptible. For example, in one patient series, ∼50% of central nervous system infections in solid organ transplant patients were caused by *C. neoformans*^6^. Treatment options for cryptococcal infection are limited and often poorly tolerated^7,8^, with case fatality rates of approximately 10% in developed settings^9,10^, 24-37% among treated HIV patients in sub-Saharan Africa^11–14^, and as high as 75% when considering all cases, treated or untreated^1^. Clinical trials have shown that combination therapy with amphotericin B and the nucleoside analog flucytosine is superior to amphotericin B combined with azole therapy^14^. However, amphotericin B can only be delivered intravenously, limiting its use in resource-poor settings, while flucytosine shortages can constrain its use^15,16^.

Among the three major classes of antifungal drugs (polyenes, azoles, and echinocandins), *C. neoformans* is exceptionally resistant to the newest (and widely used) class, the echinocandins, which target the cell wall biosynthesis enzyme β-1,3-glucan synthase; consequently, echinocandins have no clinical role in the treatment of cryptococcosis^17–20^. Since the gene encoding β-1,3-glucan synthase, *FKS1*, is reported to be essential for viability,^21^ and the *C. neoformans* enzyme is highly sensitive to inhibition by echinocandins *in vitro*,^17^ other explanations have been sought for this lack of efficacy. It has been proposed that mitochondrial sequestration limits Fks1 inhibition by the echinocandin caspofungin^22^. Loss of the lipid flippase subunit Cdc50 produces sensitivity to caspofungin but not to other echinocandins, and Cdc50 is thought to affect sensitivity in part by impacting drug uptake and/or limiting activation of calcineurin-dependent stress adaptation mechanisms^23,24^. A natural product screen identified butyrolactol A as a compound that directly targets the Cdc50-Apt1 flippase complex and increases the susceptibility of *C. neoformans* to all tested echinocandin drugs^25^. Like prior studies, this study stated that *FKS1* is essential for viability and assumed that the impact of caspofungin on *C. neoformans* growth upon Cdc50-Apt1 inhibition is due to Fks1 inhibition^25^. A recent transposon mutagenesis study classified *FKS1* as essential in *C. neoformans*^26^. We therefore sought to leverage recent advances in genome-scale editing techniques to identify potential therapeutic targets and dissect the mechanisms of drug insensitivity.

We describe here CRISPR-insert, a scalable genome-wide insertional mutagenesis strategy. We first targeted all annotated genes for mutagenesis and defined essential genes by identifying genes that could not be disrupted by insertion. We achieved this by sequencing junctions between sgRNA-encoding sequences and the genome across multiple mutant pool replicates. We used two orthogonal machine learning (ML) approaches to call essential genes, predicting 1,297 essential genes using a liberal threshold and 1,103 genes using a conservative threshold. Unexpectedly, we found that *FKS1* was absent from both sets of predicted essential genes, a result that we confirmed by generating *fks1*Δ knockout strains in multiple strain backgrounds. We performed solid-state NMR spectroscopy on intact cells, which revealed that *fks1*Δ strains lack detectable β-1,3-glucan, arguing against a cryptic, compensatory β-1,3-glucan synthase. To identify potential targets for broad-spectrum antifungal therapy development, we experimentally validated a set of genes ML predicted to be essential, whose orthologs are essential in model yeasts and that lack a human ortholog. Because combination therapy for *C. neoformans* is the clinical standard, we also developed genetic methods to quantify pairwise buffering/alleviating and synthetic/aggravating genetic interactions (GIs) on a genome scale, which, to our knowledge, have not been described in any fungal pathogens. We developed a streamlined non-essential gene library that enables “one shot” genome-scale screens. We applied this method to a panel of gene-disrupted strains to reveal GIs and found that a pathogen-specific hyaluronic acid synthase complex becomes essential in cells lacking the druggable calcineurin signaling pathway. By combining high-throughput studies with extensive experimental validation, this work identifies *bona fide* therapeutic vulnerabilities of a drug-resistant human pathogen.

## RESULTS

### Genome-wide Cas9/sgRNA-directed insertional mutagenesis approach

Motivated by prior work^27–60^, we reasoned that it should be possible to use CRISPR-Cas9 to perform genome-wide NHEJ-dependent insertional mutagenesis in *C. neoformans*. Using our optimized method^61^, we observed that transformation of a Cas9-expressing strain with an *ADE2*-sgRNA expression segment fused to a *NAT* nourseothricin resistance marker without homology arms yielded thousands of colonies on nourseothricin plates after a single electroporation (Fig S1A,B). Virtually all colonies were red (>99%), indicating disruption of *ADE2*. This combination of efficiency and specificity suggested that a pooled library approach targeting all genes was feasible. However, we first sought to determine how often two different constructs could enter and edit a single cell, as could occur with a pool and confound the analysis. We transformed cells co-expressing mCherry (mCh) and mNeonGreen (mNG) with sgRNA expression cassettes targeting either the mCh coding sequence, the mNG coding sequence, or both (Fig S1C-H). Flow cytometry of the selected transformant pool indicated specific disruption of the targeted fluorophore when only one cassette was introduced (>98%, Fig S1E). When both cassettes were transformed, a minority of cells were double mutants. Reducing the input DNA concentration reduced the frequency of double mutant cells (Fig S1F,H). In doubly transformed cells, the proportion of mCh mutants was higher than that of mNG mutants, suggesting different sgRNA efficiencies. Indeed, when we tested cassettes encoding different sgRNAs targeting either mCh or mNG, we observed similar results, but now with the proportion of mNG mutants being higher (Fig S1G,H). These data show that uptake of multiple sgRNAs can occur; we therefore calibrated electroporations to use the minimum quantity of DNA necessary to achieve the desired transformation efficiency.

We proceeded with a CRISPR-Cas9 genome-wide insertional mutagenesis scheme (CRISPR-insert) (Fig 1A). We generated a recipient plasmid pool: a Unique Molecular Identifier (UMI)-containing^62^ plasmid pool carrying the *NAT* marker, a U6 promoter, an sgRNA scaffold, and an RNA polymerase III terminator. We designed sgRNAs and selected up to ten predicted optimal sgRNAs for each gene in the genome (see Methods). This Version 1 (V1) library contained 68,792 sgRNAs targeting 6,905 genes (of 6,962 annotated nuclear genes—see Methods). We cloned an oligonucleotide pool encoding the V1 guides downstream of the U6 promoter in our plasmid pool and amplified the pool to produce linear DNA for electroporation of optimized Cas9-expressing *C. neoformans* followed by selection for nourseothricin resistance. We generated two replicate pools in each of four independently constructed Cas9-expressing strains in the KN99α strain background^63^. We isolated DNA from a portion of these pools after 36 hours of growth under selection. We then reinoculated cultures into fresh rich media (YPAD) and isolated DNA from cells after allowing for 24 further hours of growth, corresponding to ∼10 generations.

**Figure 1.**
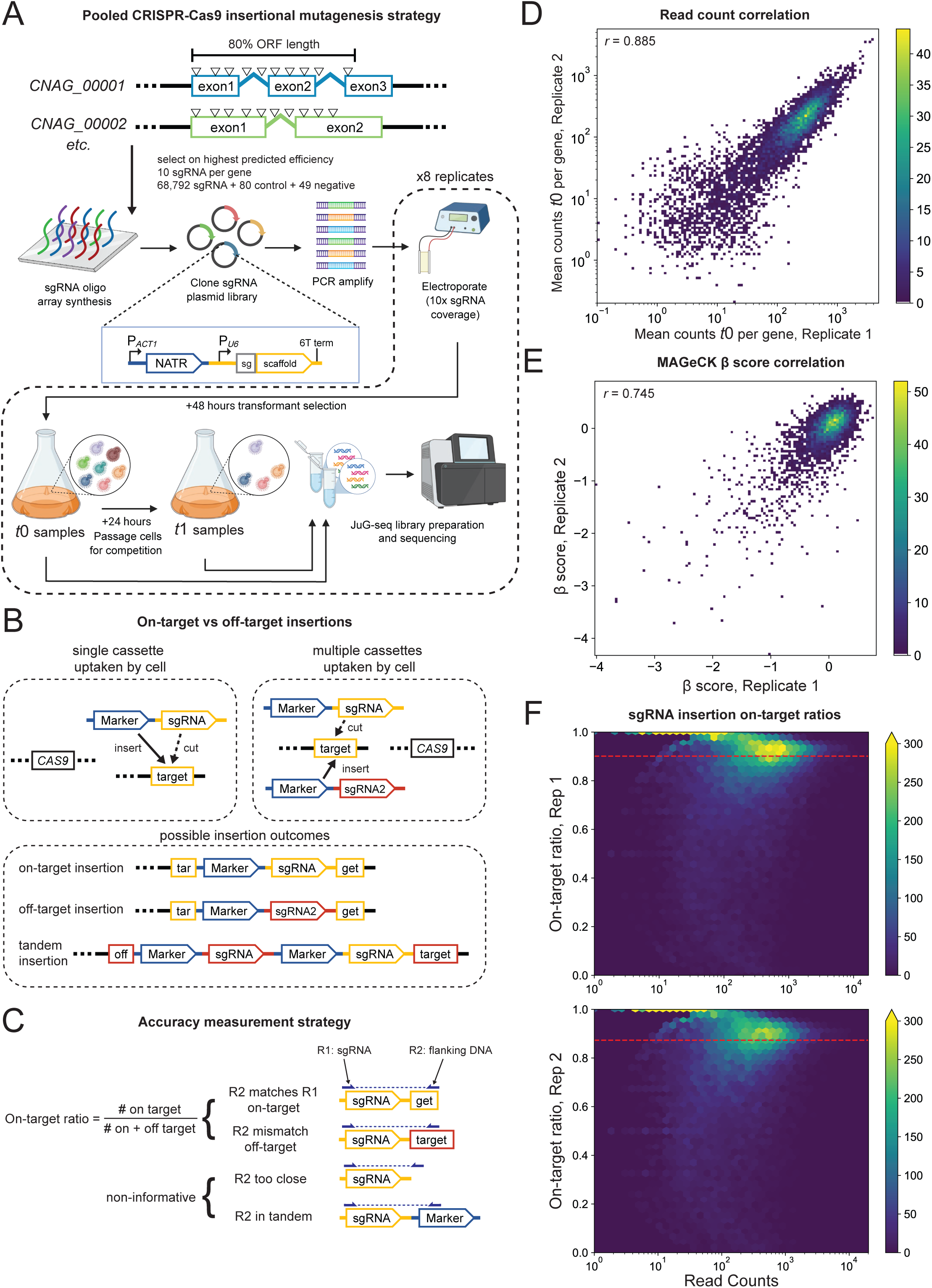
Genome-wide insertional mutagenesis with CRISPR-insert. (A) Experimental workflow of the version 1 sgRNA library covering design, transformation, assay conditions, and sequencing. (B) Expected outcome from mutagenesis occurs when an sgRNA cassette is taken up by a cell, but when multiple sgRNAs enter, a mutant may be marked by the wrong sgRNA. Possible insertion outcomes are diagrammed. (C) Simplified workflow for junction-guide sequencing (JuG-seq), showing the sgRNA and flanking junction reads. Diagram depicts classification of outcomes and defines calculation for on-target ratio. (D) Density scatterplot of mean counts per gene across all sgRNAs from two replicate mutant pools. Pearson *r* = 0.885 (E) Total counts per sgRNA from JuG-seq were analyzed using MAGeCK between initial and final timepoint samples of two replicate mutant pools. A density scatterplot of resulting β scores, analogous to LFC, is shown. Pearson *r* = 0.745. Gradient bars to the right of D and E denote density per pixel in each scatterplot. (F) Hexbin scatterplots comparing on-target rate against total read counts per sgRNA for two replicate mutant pools in CM2049. Dashed red line indicates median sgRNA on-target rate. Gradient bars to the right of each panel denote density per hex.

### JuG-seq reveals high mutagenesis accuracy

To assess mutagenesis accuracy and mutant abundance, we performed paired-end sequencing of genomic insertion junctions and the corresponding guide sequences (JuG-seq for Junction-Guide sequencing) from our mutant pools before and after 24 hours of growth in rich media. This workflow is a modification of KO-seq, which we described previously^64^ (Fig 1B,C, S2). Across all mutant pool replicates, an average of 10% of sgRNAs were absent from the mutant pool in initial timepoints, presumably due either to low sgRNA efficiency or target essentiality. On average, guides with at least one on-target read at the expected locus across initial timepoints covered 6,469 genes, of which 5,486 genes had at least five sgRNAs with at least one on-target read (Table S3). The mean counts per gene were correlated between replicates (*r* = 0.885) (Fig 1D). We used the MAGeCK-MLE statistical package to calculate fitness (β) scores from sgRNA counts for each replicate between initial and final timepoints^65^. Fitness scores from growth in rich media correlated between replicates (*r* = 0.745) (Fig 1E). For each sgRNA (read 1), we asked whether the junction (read 2) was near the expected insertion position, calculating an on-target fraction for each sgRNA as the fraction of on-target (matching) reads over all informative reads. For all identified sgRNAs with at least one on-target read in the initial timepoints of two replicate pools, median on-target fractions were high in our parental Cas9-expressing strain, CM2049 (0.90 and 0.87). On-target fraction trended even higher for more abundant sgRNAs (Fig 1F, S3).

### Machine learning predicts essential genes

Because knowledge of gene essentiality is important for defining targets for costly antifungal drug development, we reasoned that we could use our JuG-seq dataset to predict essential genes. We used two ML approaches to classify essential genes using the six highest on-target/total count ratio guides from each timepoint, which limits the impact of poorly performing guides (see Methods) (Fig 2A-B). The first approach assumed that orthologs of genes essential in other fungi would be essential in *C. neoformans*. We built a random forest (RF) classifier using ortholog conservation to label our dataset (see Methods). We considered *S. cerevisiae*, *S. pombe*, and *C. albicans* due to their well-annotated essential gene datasets. We also considered overlaps between these essential sets, under the assumption that genes essential in multiple species would be essential in *C. neoformans*. In total, we considered five label sets: one for each species alone, one for the intersection of *S. pombe* and *S. cerevisiae*, and one for the intersection of all three species.

**Figure 2.**
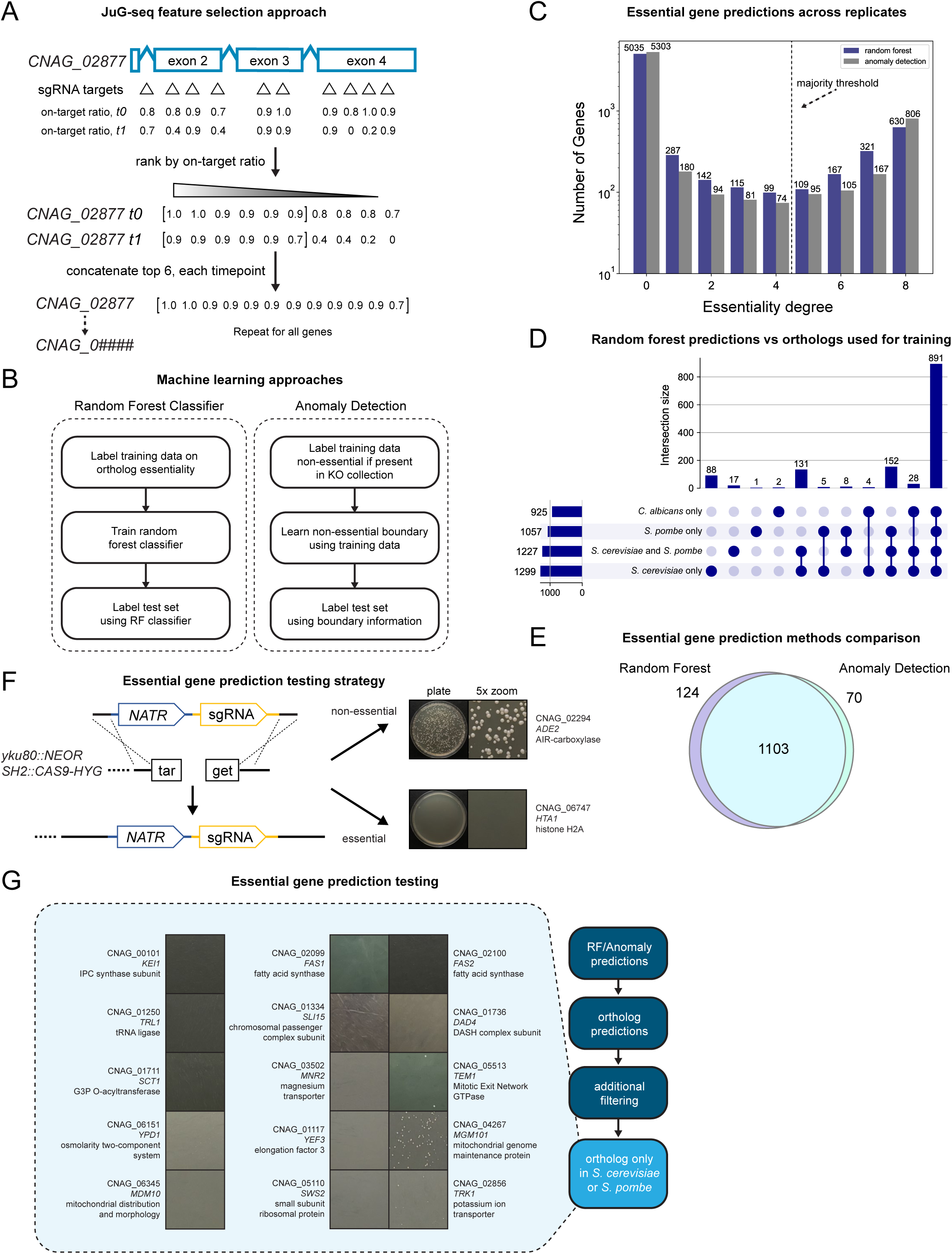
Machine learning approaches to identifying essential genes. (A) Strategy for feature selection. Top 6 on-target ratios are selected for *t_0_* and *t_1_*for each gene and concatenated for training. *CNAG_02877* was pulled as an example to illustrate feature selection. (B) Workflow for training a random forest classifier and anomaly detection classifier. (C) Bar graph showing number of replicates calling a gene essential (degree of essentiality) for both anomaly detection and random forest classifier. (D) Upset plot comparing overlap of essential calls by random forest classifier (degree > 4) using different labeled training data sets. Upset plots describe the number of elements (bar graph) that belong exclusively to the intersection of sets indicated by filled dots. (E) Venn diagram comparing overlap of essential calls by final random forest classifier and anomaly detection methods. (F) Workflow for validation of candidate genes by deletion. Transformation in a y*ku80* knockout background expressing *CAS9* will yield colonies for non-essential genes (*ADE2*) and not for essential genes (*H2A-1*). (G) Workflow for prioritization of validation targets and transformation results. Selected genes were predicted to have *S. cerevisiae* and *S. pombe* but not human orthologs. Magnified images from representative transformation plates from deletion attempts of essential gene candidates are shown.

In our second approach, we leveraged the *C. neoformans* systematic deletion collection our laboratory recently reported, which contains 4,328 validated knockout strains^64^. These genes are non-essential, as they can be deleted to produce a viable strain. We hypothesized that essential genes (and genes that are otherwise challenging to delete) could be identified as anomalies relative to the distribution of non-essential genes in an appropriate, rich-representation space derived from our JuG-seq data. Therefore, we implemented an anomaly detection (AD) algorithm that learns the boundaries of the distribution of non-essential genes in the feature space (see Methods). We posited that essential genes would be detected as anomalous and fall outside these learned boundaries. For both approaches, we used five-fold cross-validation, dividing the total available genes into five equal splits. We took each split in turn, treating it as a test set while training the classifier on the other four splits.

Table 1 shows the average and standard deviation of the evaluation metrics over these five folds for both approaches. We evaluated the effectiveness of training the RF classifier by how well our classifier recovered ortholog essentiality labels. Because the anomaly detection approach did not rely on orthologs, we used our list of non-essential genes (i.e., those present in the gene deletion collection) as a ground truth (of non-essentiality) and reported precision and recall with respect to these labels. For completeness, we extended the same metrics to the ortholog-based supervised classification experiments. The accuracy of the RF classifier was highest using the three-species overlap label set, followed by the *S. pombe* and *S. cerevisiae* intersection label set (Table 1). We note that the RF model accuracy may be inflated by the small size of the three-species label set used (Table 1). In terms of precision, recall, and F1-score, all models performed similarly, with AD performing best on F1-score and comparably on precision and recall. This is expected since the anomaly detection approach was trained specifically on the non-essentiality labels used to calculate precision and recall. We replicated our analysis for each of our eight mutant pools, obtaining eight essentiality predictions. AD classified genes slightly more consistently across pools, achieving unanimous essential or non-essential calls for the largest number of genes (Fig 2C). Surprisingly, we found substantial disagreement among essential calls in our RF classifier depending on the label set used (Fig 2D). As the *C. albicans* essentiality labels originate from Tn-seq experiments that also utilized ML training in which *S. cerevisiae* and *S. pombe* ortholog essentiality was a label, we excluded this dataset and selected the *S. cerevisiae* and *S. pombe* intersection label set as our final random forest classifier label set. Table S4 summarizes predictions for each pool.

**Table 1.**
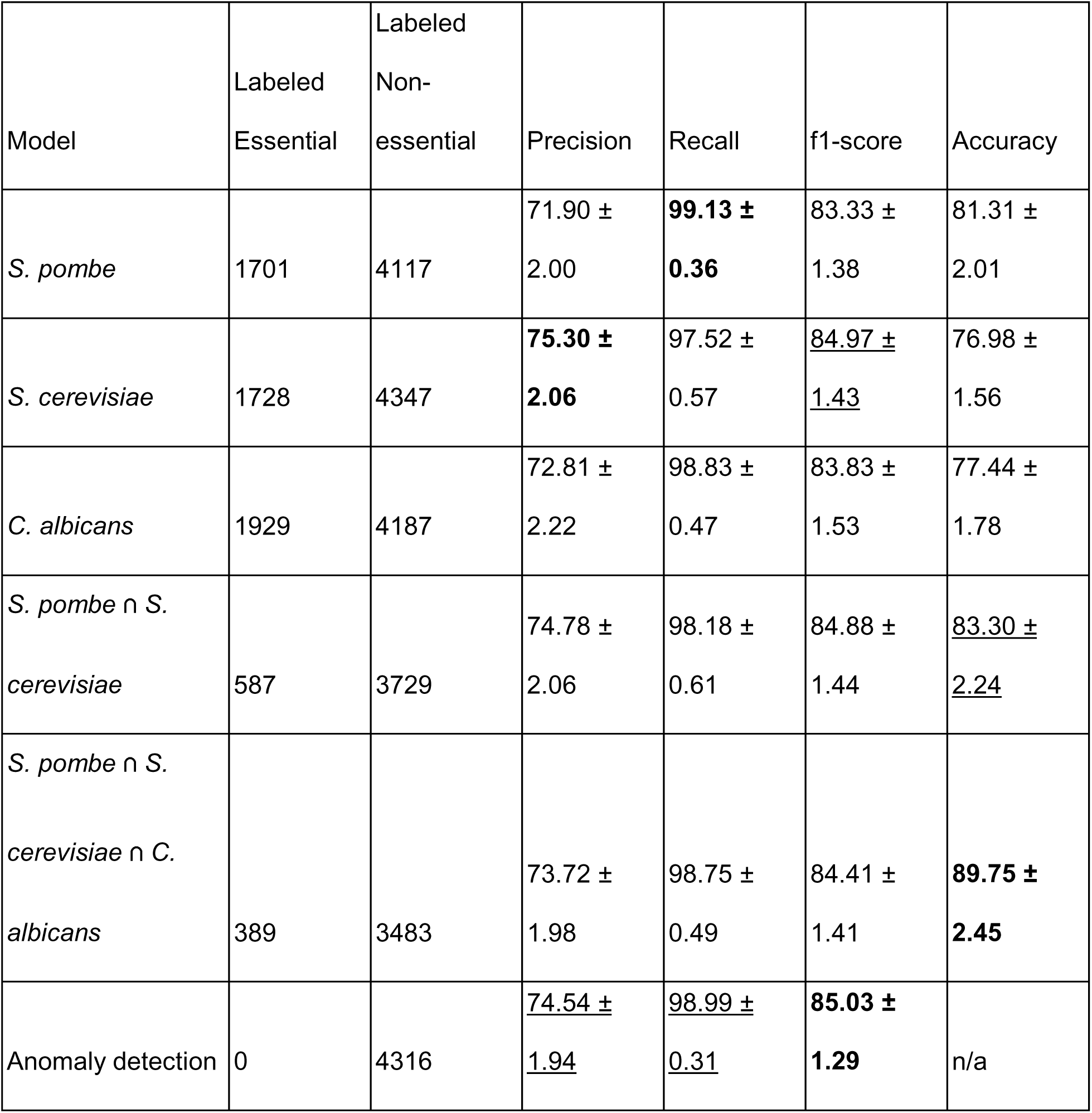
Quantitative results of ortholog-based supervised and anomaly detection approaches for essential gene predictions. Best results for each column are in bold, second-best are underlined.

| Model | Labeled Essential | Labeled Non-essential | Precision | Recall | f1-score | Accuracy |
|---|---|---|---|---|---|---|
| *S. pombe* | 1701 | 4117 | 71.90 ± 2.00 | **99.13 ± 0.36** | 83.33 ± 1.38 | 81.31 ± 2.01 |
| *S. cerevisiae* | 1728 | 4347 | **75.30 ± 2.06** | 97.52 ± 0.57 | 84.97 ± 1.43 | 76.98 ± 1.56 |
| *C. albicans* | 1929 | 4187 | 72.81 ± 2.22 | 98.83 ± 0.47 | 83.83 ± 1.53 | 77.44 ± 1.78 |
| *S. pombe* ∩ *S. cerevisiae* | 587 | 3729 | 74.78 ± 2.06 | 98.18 ± 0.61 | 84.88 ± 1.44 | 83.30 ± 2.24 |
| *S. pombe* ∩ *S. cerevisiae* ∩ *C. albicans* | 389 | 3483 | 73.72 ± 1.98 | 98.75 ± 0.49 | 84.41 ± 1.41 | **89.75 ± 2.45** |
| Anomaly detection | 0 | 4316 | 74.54 ± 1.94 | 98.99 ± 0.31 | **85.03 ± 1.29** | n/a |

Because we had eight datasets, we applied a conventional threshold based on whether the majority of pools classified a gene as essential (Fig 2C). This is a more liberal approach than requiring that a gene be classified as essential in all eight samples, each based on an independent transformation. We classified 1,227 essential genes using random forests and 1,173 using anomaly detection (Fig 2E). These sets overlap substantially, with 1,103 essential gene calls in their intersection. The union of the sets contains 1,297 genes.

### Validation of gene essentiality identifies therapeutic vulnerabilities

To identify candidate therapeutic targets, we next focused on validating genes predicted by ML to be essential. To prioritize genes, we searched the genomes of *S. cerevisiae*, *S. pombe*, and humans for orthologs of *C. neoformans* proteins predicted to be essential by ML (Fig 2G). To identify candidates for broad-spectrum antifungal drug development, we focused on 15 genes whose orthologs are essential in model yeasts but lack a detectable human ortholog based on InParanoid-DIAMOND prediction and manual literature searches (see Methods for inclusion criteria). We sought to assess the gene essentiality predictions by attempting full deletions of candidate genes in a *yku80*Δ mutant, reasoning that *bona fide* essential genes should not yield viable transformants^66^. Transformation of a Cas9-expressing *yku80*Δ mutant with a marker-guide construct targeting the non-essential gene *ADE2* yielded abundant colonies after two days of incubation on selective plates, while targeting the essential gene encoding histone H2A resulted in no colonies (Fig 2F). We used this assay to test the 15 genes described above. 13 of the 15 candidates did not produce abundant visible colonies (Fig 2G), consistent with essential roles, with the exceptions being *MGM101* and *TRK1* . In addition, we observed exceptionally rare colonies (Fig 2G) for *FAS2*, *DAD4*, and *TEM1* at a frequency about 100-fold lower than in control transformations. In each case, genotyping demonstrated deletion of the targeted gene. The rarity of these colonies suggests they may correspond to clones harboring suppressor mutations of essential genes. Suppressors of essential gene deletions have been described in *S. cerevisiae*^67,68^, where they are surprisingly frequent. Attempts at deleting *MGM101* and *TRK1* yielded abundant but small colonies, consistent with an important but non-essential role.

As we found two false-positive essential predictions, we tested 19 additional *C. neoformans* genes predicted to be essential by ML to evaluate our classifiers. Six were predicted to have essential orthologs in *S. cerevisiae* and *S. pombe* but failed subsequent manual filtering for human orthologs. All 6 did not produce abundant visible colonies, although a rare (potential suppressor-containing) colonies were confirmed for *MIS12* (Fig S4). 13 predicted essential genes are conserved in humans but lack orthologs in *S. cerevisiae* and *S. pombe*. We selected these genes for testing as they may enable studies of these human-conserved processes in a tractable yeast. At two days after transformation with a deletion marker-guide construct, we only observed abundant colonies for one gene, CNAG_04447, of 13 genes in this category (Fig S4). The others include orthologs of three human spliceosome-related factors: NKAP, ZNHIT2, and DDX41. Notably, germline mutations in DDX41 cause an acute myeloid leukemia syndrome^69^. The genes encoding a subunit of and an assembly factor for human complex I of the electron transport chain (NDUFB9 and NDUFAF5, respectively) were predicted as essential by ML, but their deletion produced abundant small colonies after three days (Fig S4). Incubation for seven days of plates from deletion attempts of six other complex I components (also classified as essential) yielded abundant but extremely slow-growing colonies, indicating an important role for complex I in *C. neoformans* growth. These data indicate that our liberal cutoff for gene essentiality prediction calls some genes essential when they are in fact non-essential, albeit important for growth (see Limitations of this study section below).

### *FKS1* is not an essential gene in *C. neoformans*

We were surprised that *FKS1* was not included in the set of 1,297 predicted essential genes (Table S4). This was puzzling since *FKS1* has been considered essential in *C. neoformans* for over a quarter century^21^ and β-1,3-glucan is both a central component of fungal cell walls and the biological ligand of the pattern recognition receptor Dectin-1 on leukocytes^70^. Although a recent study reported that *C. neoformans* growth was surprisingly resistant to transcriptional repression of *FKS1*, the authors did not take this as evidence that *FKS1* is non-essential^71^. Therefore, we revisited whether *FKS1* is essential. We recently described a strategy for near-100% efficient homology-dependent editing in *C. neoformans* using a marker-guide construct flanked by targeting homology and an NHEJ-defective *ku80* mutant strain expressing optimized Cas9^66^. Using this strategy, we found that, two days after transformation, the *FKS1* coding sequence could be readily deleted in the KN99α strain background, yielding visible colonies at a frequency comparable to a positive-control deletion of the non-essential *ADE2* gene. We genotyped *FKS1* transformants by amplifying and sequencing the locus, which confirmed complete ORF deletion and the absence of the wild-type allele. To determine whether this finding was specific to the KN99α strain background, we integrated our optimized Cas9-encoding construct into four additional strain backgrounds, A1-35-8, Bt88, Ze90-1, and FTC146-1, representing three different *C. neoformans* clades (VNIb, VNBII, VNBII, and VNBI, respectively^72^) and then inactivated the NHEJ factor Ku80. We readily deleted the *FKS1* coding sequence by homology-dependent recombination in each of these strains (Fig 3A-B).

**Figure 3.**
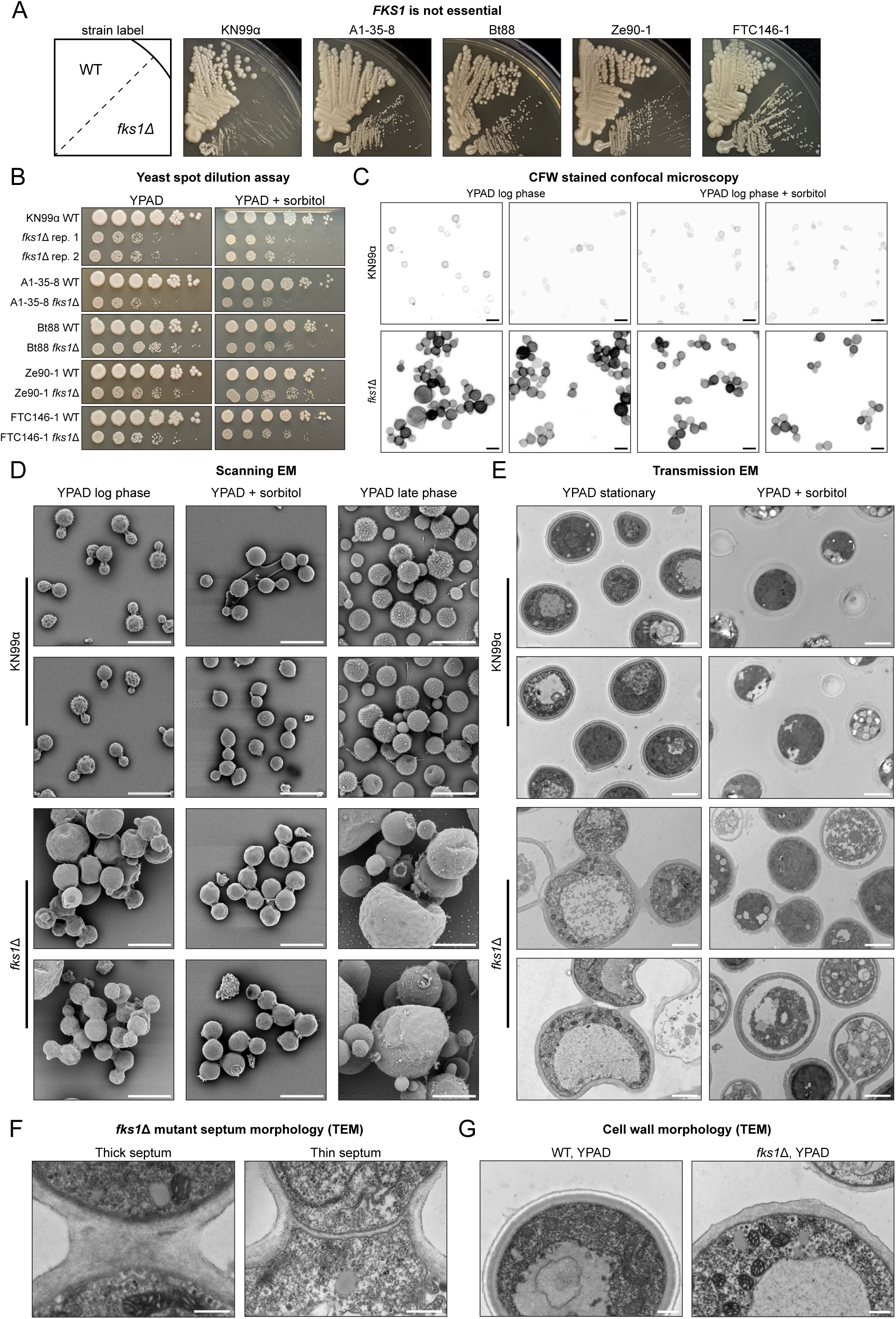
*FKS1* is not essential. (A) Wild-type isolates and *fks1*Δ mutants in isolate backgrounds patched onto YPAD media and grown for 3 days at 30°C. (B) Yeast dilution spot assay of wild-type isolates and *fks1*Δ mutants in isolate backgrounds grown for 3 days at 30°C on YPAD or YPAD supplemented with 1 M sorbitol. (C) Confocal imaging of KN99α *fks1*Δ mutant and wild-type cells stained with calcofluor white grown to log phase in YPAD and YNB with and without 1 M sorbitol supplementation. Black scale bars are 10 µm in length. (D) Scanning electron microscopy of *fks1*Δ mutant and wild-type cells grown in YPAD to stationary/late or log phase and YPAD + 1 M sorbitol to log phase. White scale bars are 10 µm in length. (E) Transmission electron microscopy of sections of *fks1* and wild-type cells grown in YPAD or YPAD + 1 M sorbitol. Scale bars are 500 nm in length. (F) TEM images showing different examples of *fks1* mutant septa phenotypes. Scale bars are 500 nm in length. (G) TEM images showing higher magnification comparison of wild-type vs mutant cell wall. Scale bars are 500 nm in length.

### *fks1*Δ strains display cell wall and morphological defects

We examined *fks1*Δ cultures microscopically using three methods: confocal imaging of live cells with the chitin-binding dye calcofluor white (CFW); scanning electron microscopy (SEM); and transmission electron microscopy (TEM) (Fig 3C-G). For this, we used the *fks1*Δ mutant in the common laboratory background strain KN99α ^63^. For the first two approaches, we examined exponentially growing cultures and late-phase, four-day liquid YPAD cultures both in the presence and absence of the osmotic stabilizer sorbitol, which typically ameliorates fungal cell wall defects. For TEM, we examined late-phase cultures. We use the description “late-phase” as it is difficult to fit the growth of *fks1*Δ mutants to a conventional growth curve.

Live-cell optical microscopy and SEM revealed a pervasive cell separation defect in exponentially growing *fks1*Δ cells (Fig 3C-D, S5A). This defect was partially suppressed by growth in sorbitol-containing medium, suggesting the phenotype is related to osmotic stress (Fig 3C-E, S5A-B). TEM confirmed a cell separation defect (Fig 3E-F). The mutant walls also lacked the even layers typical of wild-type cells (Fig 3G) and did not thicken in response to high-osmolarity medium (Fig 3E). CFW staining was higher in mutant cells, suggesting increased chitin and/or accessibility (see below).

Late-phase *fks1*Δ cells displayed considerably more aberrant morphologies. Most dramatic was the appearance of very large cells (Figs. 3C-D, S5A,C). CFW staining revealed a wrinkled phenotype of these cells, as if the cell wall was partially collapsed, an impression supported by electron microscopy (Figs. 3D-E). These data support the requirement for β-1,3-glucan in completing cell division, in maintaining cell wall strength and organization, and in responding to osmotic pressure.

### Solid-state NMR spectroscopy reveals absence of β-1,3-glucan in *fks1*Δ cells

In principle, it is possible that a cryptic β-1,3-glucan synthase exists in *C. neoformans* and compensates for the loss of *FKS1*. To test this hypothesis, we used solid-state NMR spectroscopy^73^ to examine the glycans of both the rigid and mobile phases of the cell wall. For KN99α, we selectively detected rigid cell-wall components using a two-dimensional ^13^C-^13^C correlation NMR approach, revealing prominent signals attributable to β-1,3-glucan, β-1,6-glucan, α-1,3-glucan, chitin and chitosan (Fig 4A,B, S6A). β-1,3-glucan was the predominant rigid polysaccharide, accounting for approximately 52% of the rigid fraction (Fig 4C). This occurred in two distinct polymorphic forms (type B^a^ and type B^b^), which likely reflect conformational differences (Fig 4C, S6C).

**Figure 4.**
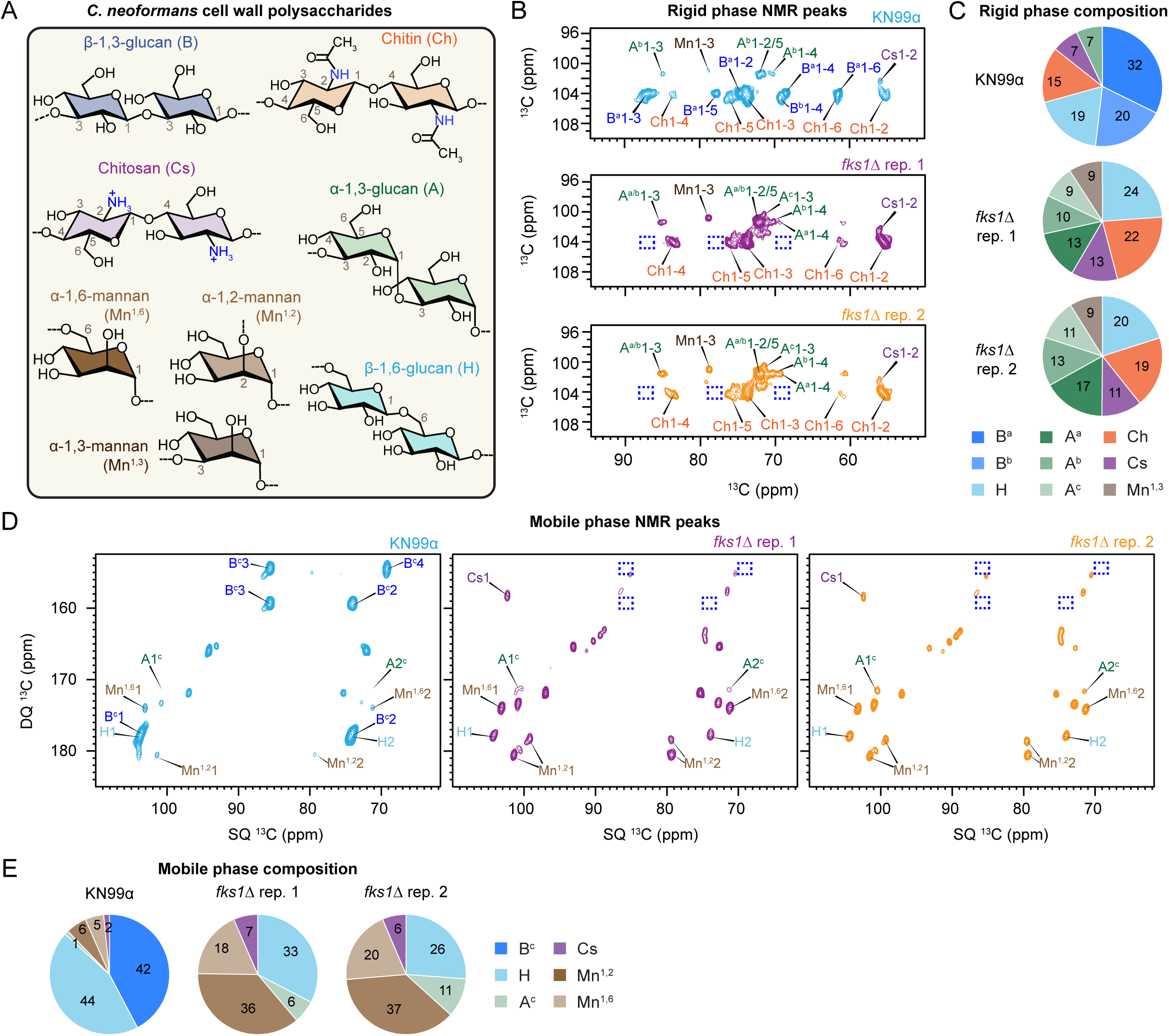
Solid state NMR analysis of cell wall glycans. (A) Simplified structural representation of polysaccharides in *Cryptococcus*, annotated with NMR abbreviations and labeled key carbon positions. (B) 2D ^13^C-^13^C CORD correlation spectra of C. neoformans cells. The absence of β-1,3-glucan signals in Fks1 mutants is highlighted with dashed blue squares. Observed cell wall carbohydrates include β-1,3-glucan (B), α-1,3-glucan (A), chitin (Ch), chitosan (Cs), and α-1,3-mannan (Mn). Two subtypes of β-1,3-glucan (B^a^ and B^b^) and three subtypes of α-1,3-glucan (A^a^, A^b^, and A^c^) are indicated by superscripts. Each cross peak represents the correlation between two unique carbons within a molecule, e.g., A^c^1-3 denotes carbon-1 to carbon-3 correlation in type-c α-1,3-glucan. (C) Molar composition of rigid polysaccharides is estimated from resolved cross-peak volumes in 2D ^13^C CORD spectra. (D) 2D ^13^C refocused J-INADEQUATE spectra obtained via direct polarization (DP) and short recycle delays for selectively detecting mobile molecules in *C. neoformans* strains. (E) Molar composition of mobile polysaccharides estimated from well-resolved peaks in 2D ^13^C DP refocused J-INADEQUATE spectra.

Remarkably, in two independently generated *fks1*Δ strains (*fks1*Δ rep. 1 and *fks1*Δ rep. 2) the signals from both forms of β-1,3-glucan were abolished (Fig 4B, blue dashed boxes). Concomitantly, α-1,3-glucan in multiple polymorphic forms^74^ substantially increased in relative abundance (Fig 4B,C). Chitin and chitosan abundance likewise displayed relative increases (Fig 4B, C). Finally, signal from α-1,3-mannan, diagnostic of capsular polysaccharide, was more readily detected, reaching approximately 9% of the rigid fraction (Fig 4B,C).

In the mobile fraction of the parental strain, β-1,6-glucan and β-1,3-glucan were the predominant polysaccharide components, accounting for approximately 44% and 42% of the mobile fraction, respectively (Fig 4D,E, S6B). These were accompanied by weaker signals from α-1,3-glucan, chitosan and mannoproteins (Fig 4D,E). Again, β-1,3-glucan signals were undetectable in both independently constructed *fks1*Δ strains, which also showed a relative reduction in β-1,6-glucan and a concomitant relative increase in α-1,3-glucan, chitosan and, most strikingly, mannoproteins (Fig 4D,E).

### One-shot genetic screens enabled by a streamlined CRISPR-insert library

Having benchmarked the CRISPR-insert V1 library, we sought to produce an optimized and streamlined sgRNA library. Such a library offers multiple advantages. Fewer sgRNAs in a library require fewer transformants and reads to maintain adequate library coverage. Selecting only known, high-performing guides improves overall library performance^75^. We removed guides corresponding to essential genes and any non-functional or low-efficiency sgRNAs that did not yield an on-target fraction of at least 0.7 (Fig 5A). Where possible, we selected 4 guides per gene. To reduce skew, we prioritized, within each gene, sgRNAs with read counts closest to the library median. The resulting Version 2 (V2) library targets 5,519 genes (covering almost all predicted non-essential genes) and contains 21,352 sgRNAs, 50 of which target neutral safe haven loci SH1 and SH2^76,77^. We ordered an oligonucleotide pool corresponding to this sgRNA library and cloned it into a plasmid pool carrying a 10N random sequence barcode downstream of the sgRNA scaffold. This 10N barcode acts as a transformant-level UMI; each transformant is uniquely marked by a combination of both UMI and sgRNA. We sequenced the pool to ascertain skew and representation (see Methods). We also changed our sequencing approach with the V2 library. Rather than implementing JuG-seq, we simplified the library preparation protocol to a single PCR, followed by sequencing the protospacer region on read 1 and the UMI on read 2 (Fig 5A). We instituted extensive washing and DNase steps to degrade library DNA bound to cell walls after electroporation (see Methods). We then used the UMIs to filter outlier reads likely originating from residual input DNA contamination or off-target events (see Methods).

**Figure 5.**
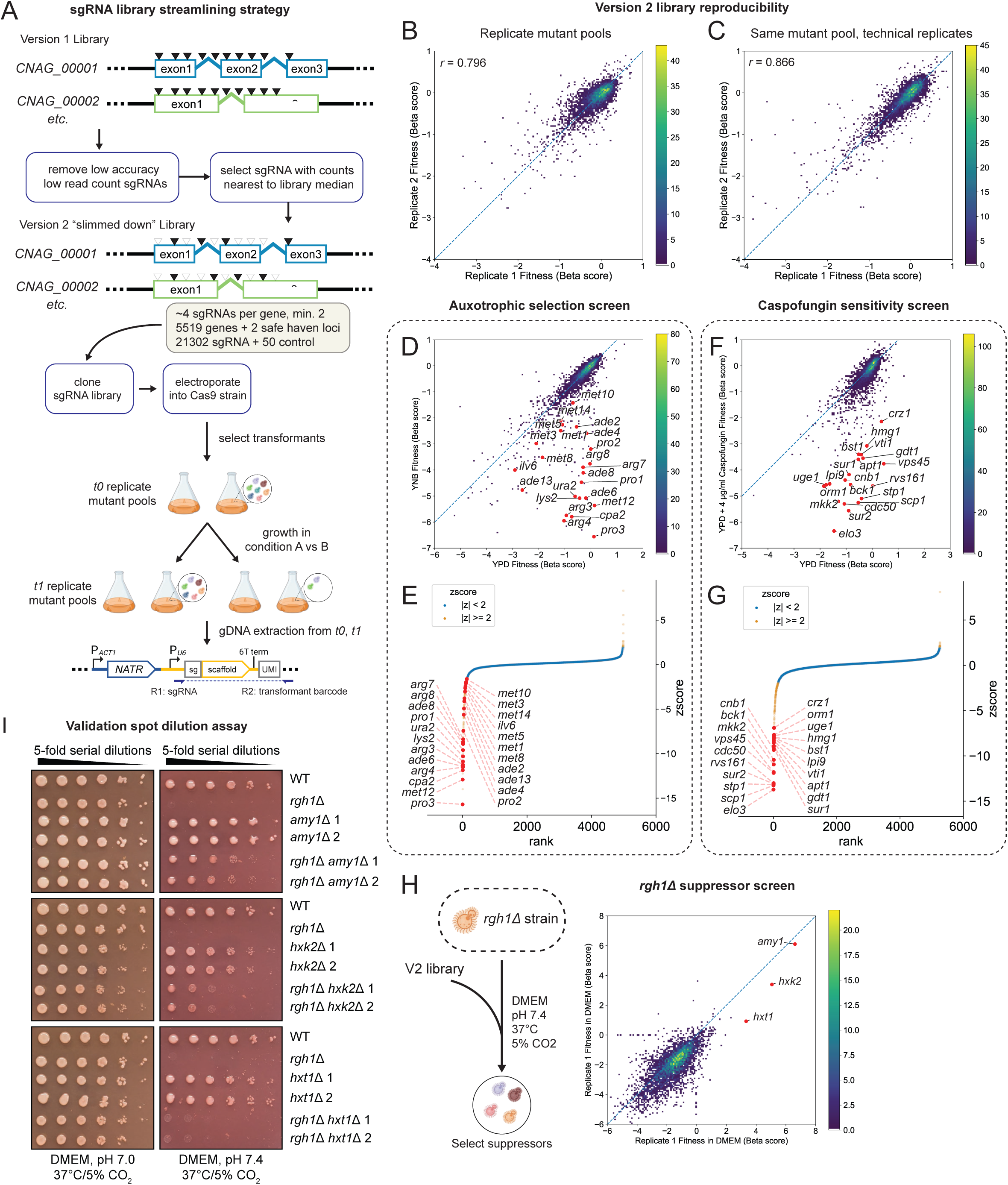
Streamlined CRISPR/Cas9 library for *C. neoformans*. (A) Strategy for design and implementation of version 2 sgRNA library based on version 1 junction sequencing data. Sequencing strategy for version 2 libraries is also simplified and reads sgRNA sequence and a transformant barcode. (B-H) For all density scatterplots, gradient bar shows density per pixel in attached plot. (B) Density scatterplot showing correlation of two independently transformed biological replicate mutant pools, Pearson *r* = 0.796. (C) Density scatterplot showing correlation of two technical replicate pools, independently grown from the same mutant pool starter culture. Pearson *r* = 0.866. (D) Density scatterplot comparing mutant fitness scores in YNB minimal media and YPAD rich media. Genes involved in production of essential metabolites are highlighted with red dots. (E) Waterfall plot on the z-score of the difference in mutant fitnesses between minimal and rich media. Values with |z-score| > 2 are highlighted in yellow. Red dots are same as in D. (F) Density scatterplot comparing mutant fitness scores in YPAD or YPAD with 4 µg/ml caspofungin. Red dots highlight 20 genes with the greatest decrease in mutant fitness in the presence of caspofungin compared to rich media. (G) Waterfall plot, same as E but for difference in fitness scores +/-caspofungin treatment. (H) Density scatterplot between two biological replicate mutant pools in an *rgh1* mutant background grown under *rgh1* restrictive conditions. Three strongest suppressors of the *rgh1* mutant are indicated with red dots. (I) Yeast spot dilution assay of *rgh1* and *rgh1* suppressor single and double mutants on DMEM buffered to either pH 7.0 or 7.4, grown in 5% CO_2_ at 37°C for 2 days.

We tested whether we could perform genetic screens using the streamlined V2 library via a single-electroporation transformation (“one shot”). We generated biological replicate pools from two independent transformations of the V2 library into our optimized Cas9-expressing strain, taking post-selection samples as an initial timepoint. We grew two technical replicates from each library in rich media. Fitness scores (as measured by β-scores produced by MAGeCK-MLE) were highly correlated between biological replicates and between technical replicates (Fig 5B,C). We tested the ability of the V2 library to detect condition-specific mutant phenotypes. We first compared pools grown in rich media vs. minimal media. Mutants in biosynthetic pathways such as those required for adenine, proline, methionine, uracil, and arginine biosynthesis had neutral fitness in rich media but were depleted in minimal media (Fig 5D,E). For some biosynthetic pathways, we did not obtain mutants whose defect in minimal media was rescued by growth in rich media, suggesting that the exogenous nutrients available in rich media are not efficiently imported. These include the histidine and tryptophan biosynthesis pathways. Indeed, our ML analysis classified all components of the histidine and tryptophan biosynthetic pathways as essential (Table S4). This contrasts with *S. cerevisiae,* where the histidine and tryptophan biosynthetic pathways are non-essential in rich media.

As a second test of our V2 library, we used our CRISPR-insert workflow to identify mutants hypersensitive to the echinocandin drug caspofungin, a screen previously performed using mutant collections^24,78,79^. Our screen identified many genes and pathways described in the literature, including the cell wall integrity (CWI) MAP kinase cascade (*BCK1*, *MKK2*, *MPK1*)^80^, the calcineurin stress-responsive signaling pathway (*CNA1*, *CNB1*, *CRZ1*)^79^, and a heterodimeric lipid flippase complex (*APT1*, *CDC50*)^24^ (Fig 5F,G). We identified additional determinants of caspofungin sensitivity, including components of the sterol-responsive SREBP pathway (*SCP1*, *STP1*), HMG-CoA reductase (the rate-limiting enzyme in sterol biosynthesis encoded by *HMG1*), sphingolipid biosynthesis and homeostasis factors (*ELO3*, *LCB2*, *ORM1*, *SUR1*, *SUR2*), and membrane trafficking factors (*RVS161*, *VTI1*).

As a third benchmark of the streamlined V2 system, we asked whether CRISPR-insert could recapitulate the identification of a suppressor of an *RGH1* mutation, a gene we recently identified as required for growth under host-like (tissue culture) conditions and virulence^64^. In a previous chemical mutagenesis screen, we identified loss-of-function mutations in the predicted α-amylase *AMY1* as suppressors of the growth defect of the *rgh1*Δ mutant under tissue culture growth conditions^64^. To test this, we generated two replicate mutant pools in an *rgh1*Δ background and grew mutant pools under restrictive tissue culture conditions. Guide sequencing of DNA extracted from the resulting cells, followed by MAGeCK analysis, identified guides targeting *AMY1* as the strongest suppressors but also identified new weak suppressors corresponding to mutations in *HXK2* and *HXT1*, which encode orthologs of hexokinase and a hexose transporter, respectively (Fig 5H,I). We validated these suppressors by generating *rgh1*Δ *hxt1*Δ and *rgh1*Δ *hxk2*Δ double deletion strains, observing that the strength of suppression in our pooled screen mirrored that observed by spot dilution assay (Fig 5I). These data show our V2 CRISPR-insert library can be used for suppressor screens with high sensitivity.

### CRISPR-insert enables quantification of buffering genetic interactions

Beyond *S. cerevisiae* and *S. pombe*, quantitative pairwise genetic interaction measurements at a genome scale have not been described in fungi, including pathogens, yet they are powerful for understanding biology^81,82^. Specifically, fully buffering interactions, in which double mutants are no more extreme than the single mutants, are often characteristic of genes that function in the same pathway, while synthetic or aggravating interactions can identify redundant pathways^83–85^.

We hypothesized that comparing the fitness scores of the 5,519 genes represented in the V2 library in wild-type versus mutant cells would allow us to measure genetic interactions. We chose a subset of the caspofungin-sensitive mutants described above as query strains. Rather than generating gene deletion mutants, we elected to preserve guide-targeting sequences and generated insertion mutants by NHEJ. A second insertion into such a mutant is anticipated to show no additional phenotype (i.e., double mutants in the same gene will display a buffering interaction). Our query strains correspond to insertion mutants of the cell wall integrity pathway (*mpk1::NEOR*, *mkk2::NEOR*), the Cdc50-Apt1 complex (*cdc50::NEOR* and *apt1::NEOR*), the calcineurin pathway (*cna1::NEOR* and *crz1::NEOR*), and the sphingolipid biosynthesis pathway (*elo3::NEOR* and *sur2::NEOR*). If buffering interactions are effectively measured, double mutants of factors in the same pathway should show among he smallest phenotypic impacts among mutants targeting the 5,519 genes represented in the library.

We transformed the V2 library into the eight genotypes described above and generated frozen stocks of the transformants. We then thawed the pools, grew them briefly, isolated DNA from a portion of the culture (*t_0_*), split the cultures into two replicates, grew each replicate in the presence or absence of caspofungin for ∼10 generations, and isolated DNA from the cells (*t_1_*) (see Methods). Following guide sequencing and data analysis, we assigned fitness scores using MAGeCK based on changes in double mutant abundance between initial and final timepoints (Fig 6A, S7). We calculated differences in mutant fitness scores between query and wild-type backgrounds under a given condition. We converted this distribution to z-scores, which correspond to our final genetic interaction scores. We set a |z|-score cutoff of 2 for interaction score significance. For buffering interactions, we focus on samples treated with caspofungin.

**Figure 6.**
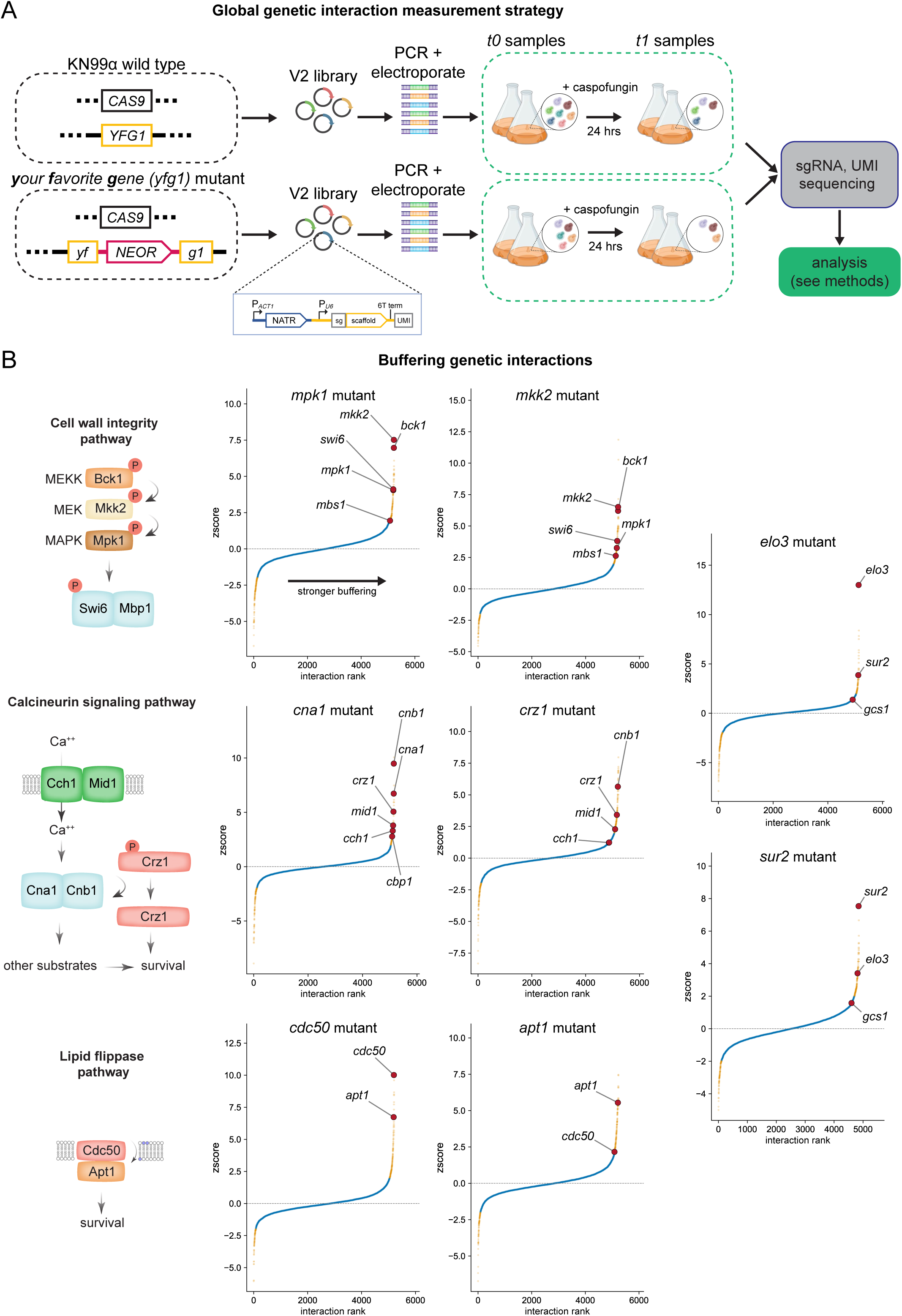
Global genetic interaction measurements in *C. neoformans* reveal pathways. (A) Strategy for genetic interaction screens. Mutant fitness is determined between *t_0_* and *t_1_* in both a wild-type and query (disruptant) background using MAGeCK. Samples for *t_1_* reflect 24 hours of growth in YPAD supplemented with EC_50_ concentrations of caspofungin. (B) Rank-order waterfall plot showing genetic interaction score (see Methods) for mutants in each query background compared to wild type in the presence of caspofungin.

Among the buffering interactions for components of the CWI MAPK pathway, we identified a subset that included other known members of the cell wall integrity pathway. For example, in the *mpk1* mutant background, *mkk2* and *bck1* mutants display strong buffering phenotypes (Fig 6B, S7). Both *mpk1* and *mkk2* show buffering interactions with mutants in orthologs of the *S. cerevisiae* SBF transcription factor complex (Mbs1-Swi6), suggesting it may be a downstream effector of this pathway in *C. neoformans*, as it is in *S. cerevisiae*^86^. We observed these buffering interactions both in the presence and absence of caspofungin; however, buffering magnitudes were greater with drug treatment (Table S5). As with the CWI pathway, we observed buffering interactions between a mutant in the calcineurin subunit-encoding gene *CNA1* and other pathway components, and similar results for the downstream transcription factor mutant *crz1*^87–91^. For example, *cna1* mutants show buffering interactions with mutants corresponding to the other calcineurin subunit Cnb1^92^, components of the upstream Cch1-Mid1 calcium channel complex^93^, calcineurin binding protein Cbp1^94^, and the Crz1 TF. We identified the same components in the *crz1* screen (Fig 6B, S7). Consistent with the literature, an *apt1* mutation is strongly buffering in a *cdc50* background, while a *cdc50* mutation is more weakly buffering in an *apt1* mutant background, suggesting that Cdc50 may form complexes with other APT family ATP-dependent flippases to influence tolerance. Finally, we observed that the sphingolipid biosynthesis enzymes Elo3 and Sur2 buffer each other in the presence of caspofungin (Fig 6B). These data demonstrate the ability of CRISPR-insert to identify buffering genetic interactions, such as those between components of functional pathways.

### CRISPR-insert enables quantification of aggravating genetic interactions

Synthetic or aggravating genetic interactions can identify redundant pathways important or essential for growth, potentially revealing targets for combination antifungal therapy. To identify such interactions, we analyzed our *t_0_* dataset for double mutants that display a growth defect relative to the single mutants (Fig 7A). Because MAGeCK produced unexpected β scores with zero-count data (as can occur with severe aggravating/synthetic interactions), we used a different statistical package, JACKS, which uses a Bayesian approach that we found suitable for this analysis (see Methods)^95^. We identified aggravating interactions for all eight genotypes tested (Table S5).

**Figure 7.**
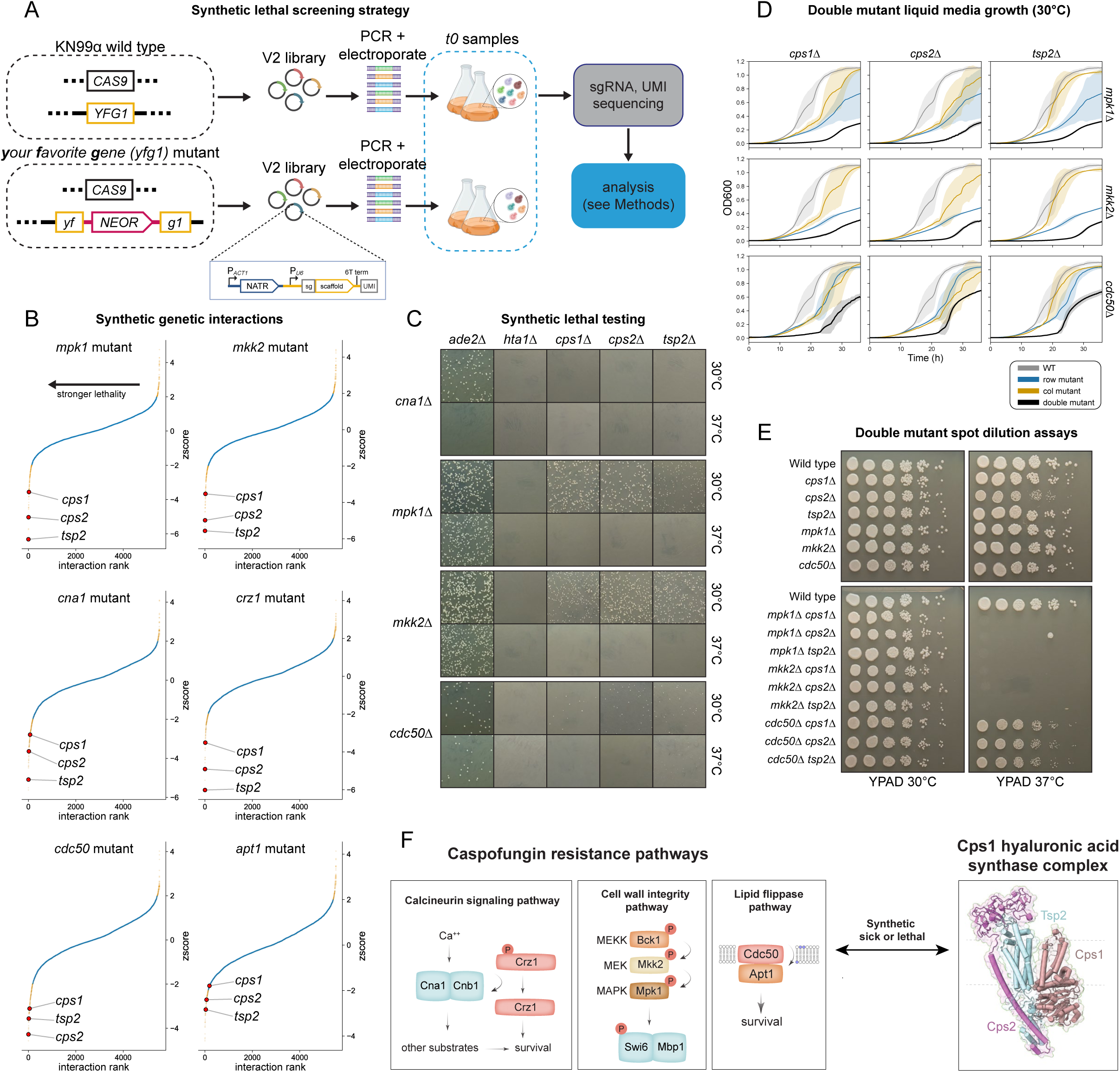
Cps1 complex displays synthetic genetic interactions with caspofungin resistance pathways. (A) Strategy for screening synthetic sick double mutants by comparing mutant pools in query (disruptant) and wild-type backgrounds using JACKS. (B) Rank-order waterfall plot showing JACKS essentiality score for mutants in each query background compared to wild type. Hyaluronic acid synthase complex genes are highlighted by large red dots. (C) Representative images from transformation plates to construct double mutant strains. Disrupted gene in parent strain is listed as the row label. Column labels refer to HAS complex component to be disrupted. *H2A-1* is included as a transformation control. Transformants were selected on YPAD + 250 ng/µL nourseothricin plates, and grown for two days at either 30 or 37°C. (D) Growth curves for HAS complex single and double mutants in *mkk2*, *mpk1*, and *cdc50* backgrounds. Row label single mutants are shown in blue; column label single mutants are show in orange; double mutants are shown in black. Shaded areas depict +/-1 SEM from 5 technical replicate wells. (E) Yeast spot dilution assays of single and double mutants grown for 2 days on YPAD at either 30 or 37°C. (F) Model diagram of synthetic sick/lethal pathways and the hyaluronic acid synthase complex.

Most strikingly, we observed that mutants in genes corresponding to the hyaluronic acid (HA) synthase complex (*CPS1*, *CPS2*, and *TSP2*) we recently reported^64^ were significantly depleted in the CWI, calcineurin, and *cdc50* pathway backgrounds, but not in mutants defective for sphingolipid biosynthesis (Fig 7B). We validated these results by constructing *cps1*Δ*, cps2*Δ, or *tsp2*Δ single and double mutants in *mpk1*Δ, *mkk2*Δ, *cna1*Δ, or *cdc50*Δ backgrounds. As with other validated essential genes, we could not obtain *cps1*Δ, *cps2*Δ, or *tsp2*Δ mutants in a *cna1*Δ background after two attempts (Fig 7C), suggesting *bona fide* synthetic lethality. We obtained viable mutants in all Cps1 complex components in *mpk1*Δ*, mkk2*Δ, and *cdc50*Δ backgrounds.

While this finding was surprising, our pooled screens examined fitness during selection in liquid media over a 48 hr time course, whereas we attempted mutant construction by plating on solid media after transformation, which can be more permissive for growth. Using these strains, we performed quantitative plate reader growth assays in liquid media (Fig 7D). For *tsp2*Δ and *cdc50*Δ, single mutants grew comparably to wild type, but *mkk2*Δ, *mpk1*Δ, *cps1*Δ, and *cps2*Δ single mutants grew more slowly than wild type. Consistent with our global measurements, double mutants of Cps1 complex components with CWI pathway components grew slowly relative to single mutants (Fig 7D). In contrast, double mutants with *cdc50*Δ showed a much longer lag phase compared to the single mutants but a minimal defect in exponential growth rates.

Because we performed all assays at 30°C, we examined growth at body temperature (37°C). Knockout transformation yielded no colonies when plated at 37°C for all tested double mutants except for *tsp2*Δ *cdc50*Δ (Fig 7C). Consistent with our transformation results, all double mutants in *mpk1*Δ and *mkk2*Δ backgrounds failed to grow at 37°C (Fig 7E). Double mutants in a *cdc50*Δ background (generated by transformation at 30°C) grew slower than wild-type at 37°C. In summary, the Cps1 complex displays synthetic genetic interactions with three pathways that control caspofungin resistance with quantitative and temperature-dependent differences. The strongest synthetic interaction was between the Cps1 complex and calcineurin. These data demonstrate the ability of CRISPR-insert to identify *bona fide* synthetic genetic interactions.

## DISCUSSION

Developing effective antifungal therapeutics is a major unmet clinical need. The only new drug class for invasive fungal infections approved in the U.S. in the last two decades has been the echinocandins, which are derived from natural products that directly inhibit β-1,3-glucan synthase, encoded by the *FKS1* gene. Unfortunately, *C. neoformans* is resistant to this class of drugs, rendering them clinically inactive. Any essential gene, particularly one that encodes a protein not found in humans, could serve as a therapeutic target. Likewise, as illustrated by the combination therapy that is the current standard of care for cryptococcal meningitis (amphotericin B and flucytosine), identifying double mutations that cause lethality offers targets for combination therapies. To identify such genes and to empower genetic studies of *C. neoformans*, we implemented genome-wide CRISPR/Cas9 and homology-independent insertion to rapidly and accurately generate and measure the fitness of mutant pools in genotypes of interest. Our CRISPR-insert pooled approach produces large, defined insertions of a marker-guide linear DNA at double-strand break sites, rather than relying on NHEJ errors to generate heterogeneous, small indels. NHEJ-mediated insertion has been used in mammalian cells to tag up to ∼5% of proteins^31^ and is widely used for genome engineering in a broad range of organisms^27–60^, but no full-genome screens involving NHEJ-mediated marker insertion to inactivate genes have been reported. Furthermore, to our knowledge, no programmed, genome-wide CRISPR/Cas9 approach has been validated in a human fungal pathogen. Below we briefly discuss key conclusions of this work.

### Application of CRISPR-insert to predict *C. neoformans* essential genes

Two ML approaches applied to data from eight replicate whole-genome CRISPR-insert screens predicted 1,297 of 6,905 genes tested as essential by either method (∼19%), with 1,103 genes (∼16%) called as essential by both methods. For *S. cerevisiae*, ∼18% of genes have been determined essential by tetrad dissection of meiotic products derived from sporulation of heterozygous knockouts^96^, while for *S. pomb*e, which has fewer genes, ∼26% of genes were reported as essential^97^. As mentioned above, a whole-genome screen applied the *S. cerevisiae* SATAY *Ac-Ds* transposon mutagenesis method to *C. neoformans* and used ML to predict 1,465 genes as essential based on ortholog essentiality (*n.b.*, experimental validation was not presented)^26^. This corresponds to 33% more genes than our low estimate and includes 108 genes present in our gene knockout collection, while our low-estimate set of 1,103 essential genes includes 18 knockouts present in our deletion collection. Because the SATAY method involves pooled growth on plates for many days followed by two liquid selection steps, lower-fitness mutants can be more easily outcompeted by higher-fitness mutants and misclassified as inviable despite training the classifier on the data labeled with orthologs essentiality^26^. In contrast, CRISPR-insert involves 1.5 day selection step and a one-day outgrowth, which limits competition. Another advantage of CRISPR-insert is that, for genetic screens, transformants from a single electroporation are selected in a single ∼100 mL liquid culture per replicate, and the method only requires modest sequencing depth, while SATAY requires hundreds of plates to generate a mutant pool, a more complex library preparation protocol, and higher sequencing depth due to the relative randomness of transposon insertions^26^. One disadvantage of CRISPR-insert is that it requires the cloning of bespoke libraries for divergent strain backgrounds due to SNPs, while SATAY retains the potential for portability across divergent strain backgrounds.

Our validation work nominates several essential genes in *C. neoformans* as potential therapeutic targets. Among these, small molecule development has been reported for some of these genes in other fungi, particularly IPC synthase (Aur1-Kei1) and fatty acid synthase (Fas1-Fas2)^98–106^. Among the remaining genes, the tRNA splicing enzyme Trl1 is an attractive target, as biochemical and functional studies have suggested for other pathogenic fungi^107,108^. Of course, essential genes with human orthologs could also be drug targets, if fungal-selective compounds can be identified. Our intersection set of 1,103 predicted essential genes provides a resource for such efforts.

### A streamlined and optimized CRISPR-insert library enables rapid genetics

By generating a streamlined and optimized V2 CRISPR-insert library, we showed that we can generate pools in which all non-essential genes are inactivated in a single electroporation and measure fitness by facile guide sequencing of DNA extracted from small liquid cultures. We benchmarked this system in three independent genetic screens, including one that identified non-essential genes whose loss produces increased sensitivity to caspofungin. This library provides a potent resource. Because NHEJ is a major repair pathway for CRISPR/Cas9 breaks in many important human fungal pathogens^109–116^, CRISPR-insert is a generalizable strategy.

### Measurement of genetic interactions in a human fungal pathogen

Despite their power to decipher biology, genome-scale genetic interaction approaches have only been developed in *S. cerevisiae* and *S. pombe* but not in other fungi, including fungal pathogens. We applied our V2 library to strains harboring insertions in genes corresponding to four pathways whose inactivation produces caspofungin sensitivity. Our quantitative analysis revealed that buffering interactions between mutants in the same pathway were readily apparent. We obtained these for the CWI MAPK pathway, the calcineurin pathway, the Cdc50-Apt1 complex, and the sphingolipid biosynthesis pathway. These findings open the possibility of large-scale GI maps to organize genes into pathways, complementing chemogenomic approaches^64,117^.

We also demonstrated the ability of CRISPR-insert to identify synthetic/aggravating genetic interactions, including synthetic lethality. We validated striking interactions between all three subunits of a Cps1 hyaluronic acid synthase complex we recently described^64^ and three pathways/complexes: the CWI MAPK cascade, the calcineurin stress-responsive signaling pathway, and the Cdc50-Apt1 lipid flippase complex. Because these pathways are linked to cell wall stress, synthetic lethality with the Cps1 complex likely reflects cell wall defects; testing this hypothesis will require detailed studies. Because Cps1 is an enzyme, it may be druggable, and since inhibitors of both calcineurin and the Cdc50-Apt1 complex have been described, developing combination therapies may be feasible. More broadly, large-scale GI screens will likely identify additional dual therapeutic vulnerabilities.

### The echinocandin target Fks1 is not essential in *C. neoformans*

*FKS1* has been thought to be essential in *C. neoformans* since it was reported thus in 1999^21^. However, our data show definitively that the gene can be deleted in multiple strain backgrounds to produce viable strains that can be cultured and analyzed. The original study was based on transforming an H99 strain background *ade2* auxotroph identified after UV mutagenesis^118^. Thus, this strain likely harbored genomic changes outside the *ADE2* gene. To test *FKS1* essentiality, the strain was transformed with a circular plasmid that harbored two inserts containing *FKS1* sequences^21^. The plasmid was designed such that a single crossover recombination with one insert would generate a full duplication of *FKS1* while recombination at the other insert would produce a disrupted allele^21^. We speculate that additional mutations in the UV-mutagenized parental strain and/or recombination bias led to the conclusion that *FKS1* is essential in *C. neoformans*. While a *C. neoformans fks1*Δ knockout has not been described previously, transcriptional repression of *FKS1* was reported to have surprisingly mild effects on growth, which in retrospect was a hint that *FKS1* might not be essential^71^. We speculate that the non-essentiality of *FKS1,* combined with other factors such as cellular drug pharmacokinetics and compensatory mechanisms, explains the remarkable resistance of *C. neoformans* to echinocandin drugs, despite the biochemical sensitivity of the *C. neoformans* enzyme to inhibition^17^. As even 100% inhibition of Fks1 in cells (which cannot occur with any drug that binds noncovalently) would not block growth, lower levels of target engagement likely have mild effects. Our analysis of *fks1*Δ cell walls by solid-state NMR spectroscopy revealed no detectable β-1,3-glucan, which argues against the presence of a cryptic β-1,3-glucan synthase that compensates for the loss of *FKS1*. In the *fks1*Δ mutant, we observe relative increases in other cell wall glycans--whether these are caused by the relative loss of the β-1,3-glucan signal versus reflecting genuine compensatory mechanisms remains to be investigated. The *fks1*Δ mutant provides a powerful tool for future genetic suppression studies.

### Limitations of this study

Gene essentiality is an operational definition based on the conditions and methods used to test it. Our transformation assay provides an operational definition, but we cannot rule out that other conditions may enable growth of a given deletion mutant. In principle, degron technology can be useful, although a lack of phenotype can reflect incomplete degradation. The acquisition of genetic suppressors of otherwise essential gene deletions, especially after prolonged incubation, is not uncommon: one study isolated bypass suppressors for 17% of essential genes queried in *S. cerevisiae*^119^. Because genetic background can affect gene essentiality, it will be important to repeat these studies across a wide range of backgrounds to distinguish background-dependent essentiality from background-independent essentiality.

## METHODS

### Culture Media

Unless otherwise specified, *Cryptococcus neoformans* was grown in YPAD media [2% w/v Bacto Peptone (#211820, BD Bacto/Difco), 2% w/v dextrose, 1% w/v yeast extract (#212720, BD Bacto/Difco), 0.04% w/v adenine] as liquid or with 2% w/v agar if solid. Strains were also grown in YNB media [0.15% w/v yeast nitrogen base powder without ammonium sulfate (#4027032, MP Biomedicals), 0.5% ammonium sulfate, 2% w/v dextrose] or with 2% w/v agar if solid. Transformed cells were grown on selective media containing, as appropriate, solid YPAD media supplemented with 125 ng/μL nourseothricin (N51200, RPI), 200 ng/μL G418 (#61-234-RG, Corning), or 300 ng/μL hygromycin B (H-270-10, Goldbio). Some transformations used *amdS* as a selectable marker - these cells were grown on solid YNB media without ammonium sulfate [0.15% w/v yeast nitrogen base powder without ammonium sulfate (#4027032, MP Biomedicals), 2% w/v dextrose] supplemented with 5 mM acetamide (A0500-100G, Sigma-Aldrich). For cloning and plasmid preparation, *Escherichia coli* was grown in LB media supplemented with 125 ng/μL carbenicillin (C-103-5, Goldbio). For cryopreservation, cells from fresh liquid cultures were kept in 0.5X YPAD or LB with 25% glycerol and stored at -70°C. DMEM for the *rgh1*Δ suppressor screen contained 13.48 g/L DMEM without sodium bicarbonate (Gibco 12800082), 50 U/mL penicillin, 50 μg/mL streptomycin (PenStrep; Gibco 15070063), 2 mM additional L-glutamine (Corning 25-005-CI), and 100 mM HEPES, adjusted to pH 7.4 with NaOH. DMEM agar plates were made similarly, except that pH was adjusted to either 7.0 or 7.4 as indicated, and 2% agar was added.

### Strain construction

Strains, primers, and plasmids used in this study are listed in Table S1. In general, we constructed all strains by electroporating the parental strain with an sgRNA expression PCR product and a selectable marker PCR product. For strains constructed by insertion, the sgRNA expression product was produced by double-joint PCR in three rounds. The first round involved two separate PCRs using the pBHM2329 template with one product made with pcnu6_R1_F and PX, and the other made with PY and sgRNA_R1_R. PX and PY are indicated for each strain in a separate column in the strain list. Round 2 did not involve primers. Round 3 used M13/F and M13/R. M13/F and M13/R were also used for amplification of the selectable marker.

For strains constructed by deletion rather than insertion, we introduced the sgRNA expression cassette and selectable marker on the same product as described^66^. Briefly, a plasmid carrying an empty guide with a *Bpl*I cut site and selectable marker was digested with *Bpl*I (ER1311, Thermo-Fisher) and the protospacer sequence was introduced using a synthesized 60-mer oligonucleotide. Cloning was performed per manufacturer’s instructions using Hifi DNA Assembly Master Mix (E2621L, New England Biolabs) and recovered by electroporation into electrocompetent DH5α *E. coli*. Plasmid sequences were confirmed by nanopore sequencing through Plasmidsaurus Whole Plasmid Sequencing (Oxford Nanopore, R10.4.1). Combined sgRNA and marker products were then PCR amplified using primers carrying homology arms as specified in the strain and primer lists.

All oligonucleotide primers were ordered from and synthesized by IDT. Unless otherwise stated, all PCR reactions used ExTaq polymerase (RR001B, Takara Bio) per manufacturer’s instructions; specific conditions are listed in Supplementary Methods.

### Electroporation

*C. neoformans* electroporations were conducted as previously described^61^. For routine electroporation, a 50 mL culture of *C. neoformans* was grown to an OD_600_ between 0.8 and 1. Cells were then collected by centrifugation (1000 x g, 5 minutes, 4°C), washed twice in 25 mL cold sterile water, and incubated in 5 mL electroporation buffer (EB) (10 mM Tris-HCl pH 7.5, 1 mM MgCl_2_, 270 mM sucrose) with 1 mM DTT (D9779-25G, Sigma-Aldrich) on ice for one hour. Cells were then collected and resuspended in 125 μL EB without DTT, which alongside residual buffer and cell volume yielded approximately 250 μL cell-buffer slurry. For each transformation, 50 μL of cell slurry was mixed by pipetting with DNA to be transformed (typically 1-2 μg of total PCR products) and transferred to a pre-chilled 0.2 cm gap Gene Pulser Electroporation cuvette (#1652086, Bio Rad). Care was taken to ensure that excess salts were not introduced to the cell slurry with the DNA to be transformed - all PCR products were column- purified and eluted in water. The cuvette was then inserted in a BTX Gemini X2 instrument and electroporated with the following settings: 500 V, 400 Ω, 250 µF. Cells were immediately resuspended in 1 mL YPAD and incubated for at least 1 hr at 30°C before plating on selective media.

The PCR product for the library transformation was amplified from 10 ng of plasmid pool DNA per 100 µL PCR reaction, using M13/F and M13/R primers and Q5 high-fidelity DNA polymerase (M0491L, New England Biolabs) per manufacturer’s instructions for 26 cycles total to avoid overamplification. Specific conditions are described in Supplementary Methods. PCR products were cleaned using NucleoSpin gel and PCR clean-up columns (#740609, Macherey-Nagel) and eluted in water. Plasmid pool cloning is described elsewhere in the methods.

### Flow cytometry

For flow cytometry of *mCherry* and *mNeonGreen* disruption transformant pools, cells were collected off transformation plates using 5 mL of YPAD and a cell spreader. Cells were then vortexed and 100 µL was then inoculated into 50 mL of YPAD + nourseothricin and grown overnight at 30°C. After growth, 1 mL of cells was taken, washed twice in PBS, diluted 1:10 in PBS, and transferred to a round-bottom FACS culture tube. Samples were then run in a BD FACSCelesta flow cytometer until at least 5,000 events passing debris and singlet gating were recorded. Events data were then analyzed using the Python FlowCal library (version 1.3.0)^120^.

### V1 screen electroporation

For V1 library electroporations, a 700 mL culture of *C. neoformans* was grown to an OD between 0.9 and 1.1. Cells were then collected by centrifugation (1000 x g, 5 minutes, 4°C), split between multiple 50 mL tubes, and washed as described above. Subsequent incubation and resuspension steps were the same as described above except all volumes were multiplied by 14. Finally, 28 μg of DNA were mixed with 3.5 mL of electrocompetent cell-buffer slurry and mixed by pipetting before being transferred to 14 pre-chilled 0.2 cm gap Gene Pulser Electroporation cuvettes (∼250 µL each). Each cuvette was subject to electroporation in a BTX Gemini X2 instrument with the following settings: 500 V, 600 Ω, 400 µF. Cells were immediately rinsed from the cuvette with 1 mL of YPAD and any leftover cells were removed by pipetting using a protein gel loading tip. Cells were then allowed to recover in a total of 70 mL YPAD in an Erlenmeyer flask for 2 hours at 30°C in an incubator with shaking. To remove excess input DNA, cells were split between 2 × 50 mL tubes and washed twice in water, once in YPAD, and twice in water again. Cells were finally resuspended in YPAD, and the entire pellet was inoculated into 1 L YPAD supplemented with 75 ng/µL nourseothricin and allowed to grow for 36 hours at 30°C in an incubator with shaking. To estimate transformation efficiency, 3×100 µL aliquots were taken at this stage and plated onto YPAD + nourseothricin plates to estimate CFUs per µL. Expected efficiency was approximately 1 × 10^5^ transformants per 50 µL electrocompetent cell-buffer slurry and we targeted at least 1 × 10^6^ transformants per replicate. After selection, 50 mL aliquots were taken either for downstream gDNA extraction or cryopreservation in 2 mL aliquots.

### V2 screen electroporation

For V2 library electroporations, a 100 mL culture of *C. neoformans* was grown to an OD between 0.9 and 1.1. Steps were the same as described above for a V1 electroporation except with routine volumes multiplied by two. This yielded 500 µL of electrocompetent cell-buffer slurry, which was mixed with 6 μg of DNA then electroporated using two cuvettes (250 µL each) and rinsed from cuvettes as described for the V1 library. Cells were then incubated in 10 mL YPAD for 2 hours in a roller drum incubator then washed as described above. Transformation efficiency was also determined by serial dilution as above, targeting 2 × 10^5^ CFUs per replicate. Cells were selected in 150 mL YPAD supplemented with 125 ng/µL nourseothricin and allowed to grow for at least 36 hours. Cells were collected as described above after we observed at least a 1 OD_600_ increase in cell density. Cryopreservation aliquots and gDNA samples were taken as above.

### sgRNA library design and cloning

We used CHOPCHOP v3 to predict efficient and specific sgRNAs for each gene in the CNA3 (RefSeq ID GCF_000149245.1) genome assembly^121^. CHOPCHOP was called with the following settings: -filterSelfCompMax 0 -scoringMethod DOENCH_2016. We then took only sgRNAs predicted to cut within the first 80% of the ORF, dropped any sgRNAs with thymine homopolymer length ≥ 4, and retained only sgRNAs with no predicted off-target sites in the *C. neoformans* genome. For each gene, we selected the 10 sgRNAs with the highest predicted efficiency based on the Doench et al. 2016 scoring metrics implemented in CHOPCHOP. We additionally designed 80 sgRNAs targeting SH1 and SH2 with the same criteria except for ORF position. Lastly, we included 49 negative control sgRNAs designed using control_guides.py from the CHOPCHOP scripts with default settings. These 68,792 sgRNAs targeting 6,905 genes and 80+49 control sgRNAs compose the V1 library.

For the V2 library we filtered the V1 sgRNAs based on the first round of JuG-seq data. We disqualified sgRNAs with no correct insertions at *t_1_*, applied a read depth cutoff at *t_0_*, then took only sgRNAs with an on-target rate > 0.7 in at least one sample. This left 35,414 sgRNAs from the original V1 library. We next ranked sgRNAs by difference to the population median read count per sgRNA. Proceeding down rank order, we discarded any sgRNA whose predicted cutsite was within 5 bp of a higher ranked sgRNA. Finally, we kept the top four sgRNAs closest to the population median for each gene, yielding 21,302 sgRNAs targeting 5,519 genes. We additionally retained 50 SH1 and SH2 control guides. Negative control guides were omitted for the V2 library. Two experiments shown in Fig 5 D,E, and H utilized a legacy library containing a subset of 19,908 sgRNAs targeting 4,977 genes.

The custom Python scripts used for calling CHOPCHOP and various filtering steps for both V1 and V2 libraries are available at github.com/madhanilab/chopchop_parallel. sgRNA library cloning

Our pooled cloning protocol was as follows: the destination vector was digested overnight, following which the digested product was run and cut out from a 0.6% agarose gel. Digested product was then purified using NucleoSpin gel and PCR clean-up columns. The insert was provided either as a PCR product or ssDNA as described in subsequent sections. NEB Hifi Assembly Master Mix was used to clone the insert into the destination vector, following which P-30 buffer exchange columns (#732-6223, Bio-Rad) were used to remove salt and dNTPs from the reaction. Products were eluted in water; then DNA was precipitated using GlycoBlue (AM9516, Thermo-Fisher), sodium acetate, and ethanol.

Approximately 500 ng of product was then electroporated into 100 µL of MegaX DH10B T1R Electrocomp cells (C640003, Invitrogen) in a single prechilled 0.1 cm gap Gene Pulser Electroporation cuvette (#1652089, Bio-Rad) in a BTX Gemini X2 instrument with the following settings: 2000 V, 200 Ω, 25 µF. A time constant within the range of 4.4 to 4.6 ms was found to be indicative of successful electroporation. Following electroporation, cells were flushed from the cuvette, and remaining cells were removed using a protein gel loading tip. The cells were then recovered in 5 mL of MegaX kit recovery media in a roller drum incubator for 1 hour at 37°C. After recovery, cells were then inoculated into a 250 mL culture of LB + 125 ng/µL carbenicillin at 30°C in an incubator with shaking overnight. The culture was also sampled immediately after recovery by plating serial dilutions onto LB + carbenicillin media to count CFUs and determine electroporation efficiency. Target CFUs exceeded insert diversity by 100-fold.

Following selection, cells in the 250 mL liquid culture were collected by centrifugation (1000 x g, 5 minutes, 4°C) for cryopreservation and plasmid preparation. Plasmid pools were extracted from bacterial cells using a HiSpeed Plasmid Midi Kit (#12643, Qiagen) with a modified protocol. Briefly, manufacturer’s instructions were followed until isopropanol precipitation. Following addition of 3.5 mL of isopropanol to the samples, 10 µL GlycoBlue was also added and samples were centrifuged at 7000 x g for 30 minutes to precipitate DNA. Pellets were washed twice in 70% ethanol and allowed to dry, after which DNA was resuspended in sterile water.

pBHM2617 is a plasmid we previously described containing the *NATR* marker and an empty sgRNA with a *Bpl*I cut site^66^. We ordered primer P03 containing homology downstream of the scaffold flanking a TruSeq adapter and a 10N UMI. P03 was cloned directly into pBHM2617 digested with *Hind*III, yielding an intermediate UMI-carrying pool. This intermediate pool was then used to clone both the V1 and V2 libraries.

The V1 library protospacer sequences flanked by homology to the U6 promoter and scaffold were ordered from Genscript as an oligo pool. The V2 library protospacer sequences flanked by homology were ordered from Agilent as a SurePrint Oligo Pool in both forward and reverse complement orientation as this was reported to reduce skew in cloning^122^. Libraries were amplified using NEBNext Q5 Ultra II Master Mix (M0544L, New England Biolabs) using primers P01 and P02 then cloned into the UMI containing pool as described above. Specific conditions and additional protocol details are described in Supplementary Methods. For quality control, the V1 library was sequenced alongside other unpublished samples in an Illumina MiSeq using a 50 cycle MiSeq Reagent Kit v2 (#MS-102-2001, Illumina) with a custom sequencing primer (P146). The plasmid library was amplified using P37 and P147 using PCR conditions as described below in “CRISPR-insert sequencing library preparation” and sequencing yielded 2.8M reads. We aligned sgRNAs to expected sgRNA sequences with an alignment rate of 76.6%, accepting only perfect matches. We detected 68,901 sgRNAs or 99.9% of the library with 31X sequencing coverage and detected on average 28.25 unique UMI sequences per sgRNA. Approximately 8.3% of the V1 plasmid pool had no insert. Skew ratio between the 90th and 10th percentiles of sgRNA abundance in the V1 plasmid pool was 2.77, demonstrating that the plasmid pool quality was well within field quality standards for sgRNA libraries^123,124^. The V2 library was sequenced alongside other unpublished samples in an Illumina NovaSeq X sequencing run without custom primers as described for other CRISPR-insert sequencing libraries yielding 26.8M reads. Perfect alignment rate was 86.55% detecting every V2 library sgRNA with 1086X sequencing coverage. Skew ratio of the V2 library was 1.51 and we detected on average 683.75 unique UMI sequences per sgRNA. Approximately 4.15% of the V2 plasmid pool had no insert. These QC data indicated that the V2 library was similarly within the expected parameters of high quality sgRNA libraries^123,124^.

### gDNA extraction

Samples for sequencing collected from screens were stored as pellets in 50 mL tubes at -70°C. For V1 library samples, pellets were treated with DNase to further remove remnant input DNA. Pellets were resuspended in 10 mL DNase I buffer (10 mM Tris-HCl, 2.5 mM MgCl_2_, 0.5 mM CaCl_2_) then digested with 30 µL DNase I (M0303L, New England Biolabs) and incubated in a water bath at 37°C for 1.5 hours. DNase I was then inactivated with 100 µL 0.5 M EDTA and heat inactivated in a 75°C water bath for 30 minutes. Cells were then washed with water before proceeding to subsequent steps.

To extract genomic DNA from samples, pellets were washed once in water then resuspended in 6 mL prewarmed CTAB lysis buffer [100 mM Tris-HCl, pH 7.4, 700 mM NaCl, 10 mM EDTA, 1% hexadecyltrimethylammonium bromide (CTAB) (H6269, Sigma-Aldrich), 1% β-mercaptoethanol (M3145, Sigma-Aldrich)]. The cell suspension was then transferred to Omni 15 mL tubes with screw caps (#19-6615-500, Omni) as the form factor of these tubes is specifically rated for subsequent beadbeating steps. Approximately 1.5 mL of 0.5 mm zirconia/silica beads (#11079105Z, BioSpec) were added to tubes, and tubes were then disrupted in an Omni Bead Ruptor Elite (#19-042E, Omni) with a 15 mL tube carriage (#17100866, Omni) with the following settings: 4 m/s, 2 x 2 minute cycles with a 90 second pause. Tubes were then incubated for 1-2 hours at 65°C. After incubation, 2 mL of 25:24:1 phenol:chloroform:isoamyl alcohol (P2069-100ML, Sigma-Aldrich) was added to the tubes and cells were then disrupted again in the Omni Bead Ruptor with the same settings. Tubes were then centrifuged at 5000 x g for 7.5 min in a fixed angle rotor at 4°C and the supernatant (approximately 7.5 mL) was removed to a fresh 15 mL tube. To precipitate DNA, 7.5 mL of isopropanol was added to the tubes and samples were centrifuged for 15 min at 3200 x g. The supernatant was then removed, pellets were washed twice with 70% ethanol, and the pellet was allowed to dry.

To further clean samples, pellets were slowly resuspended in 400 µL of TE buffer then treated with 1 µL of 10 mg/mL RNase A (EN0531, Thermo-Fisher) and incubated in a 37°C water bath for 30 minutes. Samples were then treated with 5 µL of 20 mg/mL Proteinase K (#25-530-049, Thermo-Fisher) and incubated in a 55°C heat block for 1 hour. A second 500 µL of 25:24:1 phenol:chloroform:isoamyl alcohol was added to the samples after which samples were vortexed and centrifuged for 10 min at 4°C, 20,000 x g. The supernatant was removed, then mixed by vortexing with 35 µL 3 M sodium acetate and approximately 1 mL 100% ethanol. DNA was allowed to precipitate at -70°C overnight then centrifuged for 30 minutes at 4°C, 20,000 x g. Pellets were dried and resuspended in water to yield DNA for downstream applications.

Some samples appeared cloudy or viscous at this step, suggesting polysaccharide contamination. These samples were further passed over a Zymo gDNA Clean and Concentrator column (D4065, Zymo Research) per manufacturer’s instructions.

### JuG-seq library preparation

To grow cells for analysis, we first determined CFU/µL in V1 mutant pool glycerol stocks by chipping approximately 30-50 µL of frozen cells out of the cryotubes and plating serial dilutions on YPAD plates. We then seeded 5 × 10^7^ CFUs in 500 mL of YPAD and grew cultures for 24 hours before collecting cells for gDNA extraction.

The JuG-seq library preparation workflow is diagrammed in Fig S2. Following gDNA extraction as described above, sequencing libraries were prepared using a modified KO-seq protocol^64^. Briefly, genomic DNA concentration was quantified by Qubit dsDNA BR Assay Kit (Q32853, Thermo Scientific) and 4 µg of genomic DNA in a total volume of 55 µL 1X TE pH 8.0 was transferred into a Covaris microTUBE (#520045, Covaris) and sheared using a Covaris M220 focused ultrasonicator per manufacturer’s instructions with settings for 1500 bp. Next, 50 µL of sheared product was removed from the microTUBE and DNA ends were processed using an NEBNext Ultra II End Repair/dA-Tailing Module (E7546S, New England Biolabs) per manufacturer’s instructions. P04 and P05 (carrying an 8N UMI for PCR deduplication) were annealed by slow cooling in a thermocycler as previously described^64^ and 2.5 µL was added to samples and ligated to sheared and end-prepped DNA using an NEBNext Ultra II Ligation Module (E7595S, New England Biolabs). Post-ligation, samples were size selected using Agencourt Ampure XP beads (A63881, Beckman Coulter) using 0.55x bead volume per manufacturer’s instructions. A first round of PCR was then conducted using primers P01 and P06 using NEBNext Ultra II Q5 Master Mix (M0544L, New England Biolabs) for 15 cycles. Afterwards, PCR products were cleaned using 0.55x volume of Ampure XP beads. A second round of PCR was conducted using primers P07 alongside P11-P22 carrying demultiplexing indices as described above for an additional 13 cycles. A final bead cleanup was conducted using 0.8x bead volume and resulting sequencing libraries were checked on an Agilent Tapestation 4200 using a High Sensitivity D5000 screentape (#5067-5592, Agilent). Specific conditions are listed in Supplementary Methods.

Paired-end sequencing (20 cycles Read 1, 55 cycles Read 2) was conducted using an Illumina NextSeq 550 using a 75 cycle kit (#20024906, Illumina) at the Gladstone Genomics Core.

### JuG-seq analysis

Custom Python scripts for read processing and analysis are available at github.com/madhanilab/jugseq_pipeline.

Briefly, index 1 reads containing the UMI were processed to accommodate umi-tools input requirements. Read 1 (sgRNA) and read 2 (junction) were then processed using cutadapt (v3.7)^125^ for adapter and quality trimming with options “-a GCGGATAACAATTTCACACAGG -Q 25 -m 20 --max-n 0”. The sgRNA sequence was next appended to the junction read namespace and the junction read was then aligned against a custom CNA3 genome using bowtie2 (v2.4.5) with options “--local -N 1”^126^. Alignments were then deduplicated using umi-tools (v1.1.2) with default settings^127^. We considered only alignments where the sgRNA sequence perfectly matched an expected sequence in the sgRNA library. We next calculated the distance between the junction read and the predicted breakpoint if they were on the same chromosome. We accepted alignments as “on-target” if the junction read aligned on the same chromosome and within 2 kb of the predicted breakpoint. Reads aligning to the reverse strand of the sgRNA cassette were considered too short to be informative (used as a sequencing library quality control metric) and reads aligning to the forward strand of the sgRNA cassette or the *NATR* marker were called as tandem insertions. Reads aligning elsewhere were considered off-target. Raw .fastq files have been deposited at SRA accession PRJNA1552763 and pipeline outputs are collated in Table S2.

### Essential gene classification

To generate replicate JuG-seq libraries, we first constructed 3 additional biological replicates of Cas9-expressing KN99α. We repeated the V1 library transformation in duplicate in all 3 replicate backgrounds, which in addition to the KN99α biological replicate, yielded 8 total replicate V1 library pools. We extracted gDNA and ran JuG-seq on both *t_0_* and *t_1_*samples from all libraries and considered only on-target and off-target outcomes from each sample. We next cleaned the raw data, excluding 50 genes with fewer than 10 sgRNAs per gene. Then, for each gene, we computed the on-target ratio for all sgRNAs at *t_0_*. The resulting values were sorted, and the six with the highest ratio were selected as features. The approach was repeated for *t_1_*, and the six highest ratios of correct counts were concatenated to the ones computed for *t_0_*, obtaining a final vector representation with 12 descriptors per gene. We employed the computed representation as input for the two alternative methods explored in this work.

We next constructed a set of target labels to be employed in a supervised classification framework with the aim of predicting essential genes in *C. neoformans*. Orthologs between *S. cerevisiae, S. pombe, C. albicans,* and *C. neoformans* were predicted using InParanoid-DIAMOND (v5.1) with default settings^128^. Genes with an identified ortholog in *C. neoformans* were next labeled according to published annotations. For *S. pombe* and *S. cerevisiae*, we selected all genes with Fission Yeast Phenotype Ontology annotation “inviable cell” in PomBase and Yeast Phenotype Ontology annotation “inviable” in the Saccharomyces Genome Database respectively^129–132^. For *C. albicans,* we selected all genes reported as essential by Segal and colleagues^133^. The above genes were assigned as “essential” and the remainder of genes with identified *C. neoformans* orthologs were assigned as “non-essential”. We considered five different sets of labels, one set for each ortholog species, plus two other combinations defined as follows: *S. pombe* ⋂ S. *cerevisiae* and *S. pombe* ⋂ *S. cerevisiae* ⋂ *C. albicans*. In the last two cases, where we intersected multiple orthologs to obtain target labels of essentiality, we considered a gene essential if it is essential in all the intersected orthologs, and non-essential if it is non-essential in all of the intersected orthologs. All the genes essential or non-essential in only a subset of orthologs were discarded from this analysis.

We trained a random forest classifier exploiting the 12-dimensional engineered feature vector per gene as input to predict the target labels derived in the previous step. To assess the robustness of the proposed approach, we adopted a k-fold approach with k=5: dividing the total available genes into 5 equal splits, selecting a split as the test set, and training a random forest classifier on the remaining four splits. The test set was subsequently labeled based on the classifier and the folds were rotated until all genes had been classified.

We also employed an Elliptic Envelope anomaly detector. For anomaly-detection labels, we selected all genes present in the systematic deletion collection and labeled them “non-essential”. Accordingly, we trained an anomaly detection algorithm exploiting the 12-dimensional feature vector computed from 4,316 of 4,328 presumably non-essential genes from the deletion collection. Analogously to the supervised approach, we performed a k-fold cross-validation with k=5 for training and testing the anomaly detector. We selected each fold as the test fold in turn and using each of the sets of remaining folds we fitted a multivariate Gaussian Distribution around the 12-dimensional feature vector for the labeled training data. This learned a boundary corresponding to an ellipsoid in the feature space in terms of Mahalanobis distance. Genes in the test set were accordingly classified essential if outside of the boundary and non-essential if inside.

The experiments were implemented in Python using NumPy and pandas for numerical computation and data handling. The machine-learning pipeline was developed with scikit-learn and included random forests for classification, anomaly detection using the Elliptic Envelope, and standard training and evaluation utilities. The source code and a complete specification of the package versions used in our experiments are available in our GitHub repository: https://github.com/Malga-Vision/Essential-gene-prediction.

### Essential gene candidate validation transformations

To broaden our ortholog identification, we relaxed search parameters. Orthologs between *S. cerevisiae, S. pombe, H. sapiens,* and the conservative set of 1,103 essential *C. neoformans* genes were predicted using InParanoid-DIAMOND (v5.1) with options “-score-cutoff 40 -seq-cutoff 0.1 -seg-cutoff 0.1 -conf-cutoff 0.05 -sensitivity ultra-sensitive -matrix BLOSUM62”^128^.

To identify fungal-specific essential genes that may function as potential vulnerabilities, the conservative set of 1,103 essential *C. neoformans* genes was filtered on the following criteria. First, we excluded any genes with a predicted human ortholog. Second, we retained only genes with orthologs in both *S. cerevisiae* and *S. pombe*. Third, we excluded any genes where the Alliance of Genomic Resources predicted a human ortholog for an *S. cerevisiae* gene^134^. Fourth, we performed a manual literature search using *S. cerevisiae* and *S. pombe* gene names to filter out any with a human ortholog. Fifth, we filtered out any genes with potentially extracellularly rescuable mutant phenotypes (e.g. vitamin or amino acid biosynthetic pathways).

To identify *C. neoformans* unique genes with potential human orthology, we used the InParanoid predictions to filter out genes with orthologs in either *S. cerevisiae* or *S. pombe* and retained genes with predicted human orthologs. We then performed a manual literature search to remove genes with potential orthologs in either *S. cerevisiae* or *S. pombe*.

We used a slightly altered strain construction protocol for validation of essential genes. We disrupted *YKU80* with an *NEOR* marker in CM2049 to yield CM2451. We then cloned sgRNAs into pBHM2644 and PCR-amplified deletion constructs as described under “Strain Construction”. CM2451 was then transformed with 250 ng of the relevant PCR product for each candidate in at least two replicate transformations for each gene. Transformants were plated and selected on YPAD medium + 250 ng/µL nourseothricin, and plates were imaged after 2 and 7 days. Increased nourseothricin concentration was used to eliminate spontaneously arising nourseothricin resistant *C. neoformans* colonies.

### *FKS1* deletion in isolates

The *C. neoformans* isolates used in this study were obtained from Dr. Joseph Heitman (Duke University) (Table S1)^72^. We authenticated strains by sequencing of diagnostic genomic fragments. To facilitate CRISPR-mediated genetic manipulation, strains were first engineered to express Cas9 by integration of the Cas9 construct at the SH2 locus. *YKU80* was subsequently disrupted by insertion of a neomycin-resistance (*NEO*) cassette, generating strains with the genotype *SH2::CnoCAS9-HYG ku80::NEO*.

For deletion of *FKS1*, sgRNAs were selected according to isolate background. Guide 1 (5′-CATTCCGAATACTACCAGTG-3′) was used for KN99α, A1-35-8, Ze90-1 and FTC146-1, whereas guide 2 (5′-CCTCCCTCAGATTACCACGG-3′) was used for Bt88. Transformation constructs were the same as described for essential gene validation. Cells were transformed according to the transformation protocol described above using 250 ng DNA and plated onto YPAD + NAT media. Plates were incubated at 30°C for 3 days followed by 2 days at room temperature. Colonies recovered following *FKS1* targeting were inoculated into YPAD broth and grown at 30°C for 48-72 h. Genomic DNA was then extracted, and deletion of *FKS1* was confirmed by PCR genotyping.

### CRISPR-insert screens

To assay mutant abundance in V2 mutant pools, libraries were recovered from frozen library glycerol stocks. CFU/µL in V2 mutant pool glycerol stocks was determined by serial dilution as described for V1 JuG-seq. At least 2 × 10^7^ CFUs were inoculated from frozen stocks into 100 mL of YPAD and grown for 16 hours at 30°C with shaking. This culture was retained as the initial timepoint for V2 screens in all backgrounds except for *rgh1*. Subsequently, a fresh culture of 100 mL of the relevant medium for each screen was inoculated at an OD_600_ of 0.02 from the initial timepoint culture and grown for 24 hours.

For caspofungin screens, backgrounds were treated at approximately the EC_50_ caspofungin concentration such that by 24 hours treated pools reached half the growth observed in the untreated condition. To determine this concentration, multiple pools were grown in incrementally decreasing concentrations of caspofungin and pools exhibiting half the growth of the untreated condition were retained.

To identify suppressors of *rgh1*, we inoculated 2 × 10^7^ CFUs from frozen library glycerol stocks of our duplicate libraries directly into DMEM, pH 7.4, and cultured with shaking at 200 rpm at 37°C/5% CO_2_ for 4 days. Cultures were then pelleted, inoculated into 100 mL YPAD cultures at an initial OD_600_ of 0.1, and outgrown in YPAD for harvesting and gDNA isolation as described above. We used the frozen library stocks as the initial timepoint reference for these cultures.

### CRISPR-insert sequencing library preparation

Following gDNA extraction, genomic DNA concentration was quantified by Qubit and 500 ng of gDNA (roughly equivalent in mass to 2.5 × 10^7^ *C. neoformans* genomes) was used as a template for PCR amplification. Using NEBNext Ultra II Q5 Master Mix, sgRNA and UMI sequences were PCR amplified from gDNA samples with custom primers carrying Illumina adapter and index sequences on overhangs. Libraries were single-indexed as P23-31 carry a heterogeneity spacer to allow use of TruSeq adapters for Read 1, but do not carry an i5 index. Legacy libraries were amplified using P147 without a heterogeneity spacer, necessitating the use of a custom sequencing primer. P32-145 anneal to the TruSeq adapter downstream of the scaffold and carry indices for demultiplexing. Specific PCR conditions are listed in Supplementary Methods. PCR products were cleaned using Monarch Spin PCR and DNA Cleanup Kits (T1130L, New England Biolabs) and resulting products were quantified by Qubit. Samples were then pooled and checked on an Agilent Tapestation 4200 using a High Sensitivity D1000 screentape (#5067-5584, Agilent) before sequencing.

Paired-end sequencing (2 x 50 cycles) was conducted using an Illumina NovaSeq X at the UCSF Center for Advanced Technologies or run on a NextSeq 500 using a 75 cycle NextSeq 500/550 High Output v2.5 kit (#20024906, Illumina) with a custom sequencing primer (P146) at the Gladstone Genomics Core for legacy V2 libraries. All such sequencing runs used a 5% PhiX sample spike-in.

### CRISPR-insert analysis

Custom Python scripts for CRISPR-insert read processing and analysis are available at github.com/madhanilab/guideseq_pipeline.

For CRISPR-insert sequencing, read 1 carried the sgRNA sequence and read 2 carried the transformant UMI. Specifically, when the sgRNA plasmid libraries were cloned, sgRNA sequences were cloned into a pool of UMIs downstream of the scaffold, yielding multiple UMIs per sgRNA sequence. Subsequently, as there are likely fewer transformants for a given sgRNA than there are possible sgRNA-UMI combinations, each transformant was assumed to carry a unique sgRNA and UMI combination.

Reads were first processed using cutadapt 4.2^125^ for adapter, quality, and size trimming with options “-q 25 -e 0.2 -m 20:10 -M 31: --max-n 0 -g acaccggcagggtatactgttg”. The sgRNA and UMI sequences were then extracted and the sgRNA was checked against the expected sequences allowing for no mismatches. A counts dictionary was built for each UMI for each sgRNA for each gene targeted. The umi-tools (v1.1.2) CLI method UMIClusterer was then used with “cluster_method = directional” to collapse UMI counts onto the highest count parent^127^. We calculated a counts threshold where given *n* total counts for a given sgRNA, threshold = max(floor(log2(*n*+1)-2), 2) to computationally remove input DNA contamination by pruning any UMIs and associated counts beow this threshold.

We next used the UMI sequence to filter jackpotting events from *t_0_*samples. We binned sgRNAs by the number (*m*) of distinct UMIs. For each bin, we calculated IQR of log-normalized counts and set the upper threshold at Q3 + (0.5 + 0.25\**m*)*IQR. If a given sgRNA had more counts than this ceiling, then those counts corresponded to an implausibly small number of transformants (UMIs), i.e. resulting from jackpotting events. We zeroed counts from any such sgRNAs and for each library dropped any sgRNAs with fewer than 10 counts in either initial timepoint replicate. We considered only genes for which we had counts from 2 or more sgRNAs, and formatted the data for MAGeCK input, comparing change in abundance between *t_0_* and *t_1_*samples with or without drug as appropriate. We then used mageck mle with options “--norm-method total --no-permutation-by-group” to calculate β scores reflecting change in abundance of each gene between timepoints^135^. Given β_query_ and β_WT_ we defined the genetic interaction score for a given condition as GIS = zscore(β_query_ - β_WT_). Collated beta scores and genetic interaction scores are reported in Table S5 under the “betas” and “interaction_scores” tabs.

To call synthetic sick double mutants, we used JACKS (v0.2) comparing wild-type and query initial timepoint samples^136^. We reformatted filtered reads from timepoint 0 samples as above and called JACKS with default settings specifying SH1- and SH2-targeting guides as control guides for *p*-value calculation. Collated essentiality scores and p-values are reported in Table S5 under the “JACKS” tab.

### Scanning electron microscopy

Samples were grown under desired conditions then washed twice in PBS and fixed overnight at 4°C in 2.5% glutaraldehyde, 2% paraformaldehyde in 0.1 M sodium phosphate buffer pH 7.4. After fixation, cells were washed once with and stored in 1x PBS at 4°C. To prepare samples for microscopy, 12 mm coverslips were plasma cleaned then treated with poly-L-lysine for 30 minutes before being rinsed in clean sterile water. Coverslips were placed in a 24-well polypropylene plate and fixed cells were then allowed to adhere to coverslips for 1 hour at 4°C without agitation. Coverslips were then gently washed twice by pipetting 1 mL sterile water into the corner of the well containing the coverslip. Samples were then sequentially dehydrated in 25%, 50%, 75%, 90%, and 100% acetone. Coverslips were then mounted in a Leica EM CPD300 and critical point dried using liquid carbon dioxide. Samples were then mounted on aluminum stubs using double-sided adhesive carbon tabs and sputter-coated with 5 nm iridium in a Leica EM ACE600 high vacuum sputter coater. Samples were then imaged at 5 kV in a Phenom Pharos 2 desktop scanning electron microscope (Thermo Fisher Scientific). Brightness and contrast adjustments as well as cropping were done in ImageJ^137^.

### Spot dilution assays

Strains to be spotted were grown overnight in liquid YPAD cultures. Cells were then counted using an Invitrogen Countess cell counting chamber slide (C10228, Fisher Scientific) and a Countess II (Invitrogen). Cells were then normalized to 3 × 10^6^ cells per mL. As *fks1*Δ mutants are clumpy, a best estimate of the average number of cells per mL was taken from multiple slides. Five-fold serial dilutions were taken from this culture for a total of six dilutions. Three µL of each dilution was spotted onto the relevant solid media with a multichannel pipette. Cells were then grown for the number of days indicated in the figure legends at 30°C or the indicated temperature and spot dilutions were photographed.

### Growth assays

Strains to be assayed were grown overnight in YPAD, then diluted to OD 0.01 in fresh YPAD. 200 µL of each dilution was then added to 5 technical replicate wells of a 96-well untreated flat-bottom plate (#351172, Corning). Blank wells were filled with 200 µL of YPAD along the left and right edges of the plate and covered with a Breathe-Easy polyurethane film (#9123-6100, USA Scientific). Cells were then grown for 36 hours and OD_600_ was measured every 15 minutes in a Tecan Infinite 200 Pro. Growth settings were as follows: 30°C, 4 mm amplitude orbital shaking. Growth curve data were then graphed using a Python script generated using Claude Code, available at github.com/madhanilab/growthcurve_claude.

### Confocal microscopy

*C. neoformans* strains were grown from frozen glycerol stocks on YPAD plates at 30°C. Single colonies were inoculated in 5 mL YPAD or YNB (0.67% yeast nitrogen base without amino acids (Difco catalog #291940) containing 2% dextrose) +/- 1 M sorbitol and grown for four days at 30°C with shaking (230 rpm). Cells were collected by centrifugation and washed twice with sterile 1X PBS. 10^7^ cells were stained with 100 μg/mL calcofluor white in PBS for 10 minutes at room temperature while rotating and washed twice with sterile 1X PBS. Images acquired with a Zeiss LSM880 confocal microscope are shown as sum projections of Z-stacks. Brightness and contrast were adjusted in Fiji consistently within each experiment^137^.

### Transmission electron microscopy

Single colonies of *C. neoformans* strains were inoculated in 25 mL YPAD +/- 1 M sorbitol and grown for four days at 30°C with shaking (230 rpm) and fixed with 2% paraformaldehyde/2.5% glutaraldehyde (Ted Pella Inc., Redding, CA). Electron Microscopy was performed as described previously elsewhere^138^.

### Preparation of ^13^C,^15^N-labeled fungal material

*C. neoformans* strains were inoculated from single colonies into 25 mL 1X YNB with 1.5% ammonium sulfate and 2.1% ^13^C-glucose and grown for 4 days at 30°C with shaking (230 rpm). For solid-state NMR characterization, 35-45 mg of natively hydrated whole-cell material was packed into a 3.2-mm magic-angle spinning (MAS) rotor (HZ16916, Cortecnet).

### 13C solid-state NMR experiments

High-resolution 1D and 2D solid-state NMR experiments were conducted at 800 MHz (18.8 Tesla) at Michigan State University. All ^13^C-detection experiments were performed using a 3.2 mm HCN probe at 15 kHz MAS, with ambient temperatures between 283 K and 298 K. ^13^C chemical shifts were externally referenced by calibrating the adamantane CH_2_ peak to 38.48 ppm, and the resulting spectral reference (sr) value was applied to fungal spectra. Unless otherwise specified, typical radiofrequency field strengths were 83-100 kHz for ^1^H decoupling, 62.5 kHz for ^1^H hard pulses, and 50-62.5 kHz for ^13^C.

1D ^13^C spectra were acquired using various polarization methods to probe molecular dynamics. Rigid components were detected via dipolar-based ^13^C cross-polarization (CP) with a 1-ms contact time, adhering to Hartmann-Hahn match conditions of 62.5 kHz for ^13^C and ^1^H. For quantitative analysis, 1D ^13^C direct polarization (DP) experiments were conducted with a long recycle delay of 35 s, ensuring full longitudinal relaxation before the next scan. Additionally, a shorter recycle delay of 2 s in the same ^13^C DP experiment enabled selective detection of mobile molecules with fast ^13^C-T_1_ relaxation.

To facilitate resonance assignment, 2D ^13^C-^13^C correlation experiment was conducted using a 53-ms CORD (combined R2_n_^v^-driven) mixing period, which revealed intramolecular cross-peaks between carbon sites within each molecule. Additionally, 2D ^13^C refocused J-INADEQUATE experiments were conducted using either CP or DP (1.5 s recycle delay) to probe molecular domains with differing mobility. This experiment correlated double-quantum (DQ) chemical shifts with two corresponding single-quantum (SQ) shifts, generating asymmetric spectra that enabled efficient through-bond carbon-connectivity tracking. To optimize carbohydrate detection, the τ period (out of four) was set to 2.3 ms. All 1D experiments, as well as 2D CORD experiments, were conducted for all *C. neoformans* strains. Data acquisition was conducted using Topspin 3.5, and spectral analysis was performed in Topspin 4.0.8.

Molecular composition analysis was conducted by selecting only well-resolved signals in 2D ^13^C CORD spectra for rigid components and ^13^C DP refocused J-INADEQUATE spectra for mobile molecules. Peak volumes were quantified using the integration function in Bruker Topspin, with quantification based on the mean of resolved signals. Relative polysaccharide abundance was determined by normalizing the sum of integrals with their respective counts. The standard error for each polysaccharide was calculated by dividing the standard deviation of integrated peak volumes by the total cross-peak count. The overall standard error was computed as the square root of the sum of squared errors for each polysaccharide. The percentage error was determined by normalizing the standard error with the average integrated peak volume and adjusting for each polysaccharide’s relative abundance.

## ACKNOWLEDGEMENTS

We thank Irene Beusch and Stefan Oberlin for their extensive advice on methodology, and Sam Lord and Natalie Petek for extensive instruction on scanning electron microscopy and Dyche Mullins for use of the instrument. We thank Mylinh Bernardi of the Gladstone Genomics Core for their assistance with NextSeq 550 sequencing.

## AUTHOR CONTRIBUTIONS

M.Y.H., P.H., and K.C. constructed all strains used in this work unless otherwise noted.

P.H. and K.C. conducted validation experiments shown in Figure 2. P.H. constructed the *fks1*Δ mutants and conducted SEM experiments shown in Figure 3. K.C., supervised by M.Y.H., conducted experiments shown in Figure 7. M.Y.H. and H.D.M. developed CRISPR-insert. M.Y.H. with assistance from P.H. and K.C. designed and cloned sgRNA libraries, conducted all screens/NGS, and analyzed the data unless otherwise noted. M.J.B. developed and M.Y.H. modified the JuG-seq workflow shown in Figure S2. M.C. and V.P.P. developed machine learning classification pipelines and analyzed data for prediction of essential genes shown in Figure 2 and Table 1. A.L.W. and M.J.B. constructed strains and performed the *rgh1*Δ suppressor screen and spot dilution assay shown in Figure 5. L.L. and V.N. constructed *fks1*Δ mutants in isolates shown in Figure 3A-B. S.L.B. supervised by T.L.D. conducted confocal microscopy, conducted TEM, and analyzed the data shown in Figure 3 and Figure S5. A.A., supervised by T.W., conducted all NMR experiments and analyzed the data shown in Figure 4 and Figure S6. M.Y.H. and H.D.M. drafted and edited the paper with input from other authors. H.D.M. oversaw the work.

## AI DISCLOSURE

Claude Code (Sonnet 4.6, 5.0; Opus 5.0) was used for assistance in writing Python scripts for figure generation. All code for figure generation was reviewed by M.Y.H. before use. AI assistance was not used in code written for data processing of CRISPR-insert or JuG-seq data. Claude Opus was used in proofreading, but not in writing of the manuscript.

## SUPPLEMENTARY FIGURE LEGENDS

**Supplementary Figure S1. Efficient target disruption by homology-free insertional mutagenesis.** (A) Strategy for insertional mutagenesis without homology arms. (B) Transformation plates from electroporation of a construct carrying *NATR* marker and ADE2-sgRNA or marker alone. (C) Experimental setup for disruption of fluorophore expression in an *mCherry* and *mNeonGreen* expressing strain with one or multiple sgRNA expressing constructs. (D-G) Density scatterplots overlaid with contour plots of flow cytometry data showing mCherry and mNeonGreen signals from transformant pools. Gradient bars indicate density per pixel. (D) Flow cytometry of parent double fluorescent strain. (E) Flow cytometry of transformant pool from fluorescent parental strain transformed with either sgRNAs targeting *mCherry* or *mNeonGreen*. (F) Flow cytometry of transformant pool from fluorescent parental strain transformed with both sgRNAs across decreasing total input DNA concentrations. (G) Same as F with alternate *mCherry* or *mNeonGreen* targeting sgRNAs. (H) Stacked bar graph of mean percentage of mutant outcomes from transformant pools using different total quantities of DNA construct. Error bars are +/- SEM; horizontal lines above data denote significant difference in double mutant frequency between samples (t-test, **, p < 0.01).

**Supplementary Figure S2. Workflow for KO-seq/JuG-seq.** (A) Diagram showing steps in preparation of JuG-seq / KO-seq sequencing libraries. (B) Workflow for computational analysis of JuG-seq data.

**Supplementary Figure S3. Additional replicate accuracy data.** (A) Hexbin scatterplots comparing on-target rate against total read counts per sgRNA. Dashed red line indicates median sgRNA on-target rate. Gradient bars indicate datapoints per hexbin. (B) Violin plots showing on-target rate for all *t_0_*and *t_1_* samples.

**Supplementary Figure S4. Additional testing of essential gene candidates.** Representative magnified images from transformation plates from deletion attempts of indicated essential gene candidates. Workflow for prioritization of validation targets is reproduced as from Fig 2G. 6 genes predicted to have only yeast orthologs but did not pass manual literature filters are shown to the left. Genes predicted to have human but not *S. cerevisiae* or *S. pombe* orthologs are shown to the right.

**Supplementary Figure S5. Additional images and quantification of *fks1***Δ **mutant phenotypes.** (A) Confocal imaging of *fks1* mutants and wild-type cells stained with calcofluor white grown in YPAD and YNB to late phase with and without 1 M sorbitol supplementation. Black scale bars are 10 µm in length. (B) Percentage of cells in the indicated conditions that are singlets, doublets, or in a clump of three or more cells. Between 80-430 cells were measured for each condition. (C) Size of cells in the indicated conditions. Measurements represent the average of the length of the longest diameter and the diameter perpendicular to it. At least 100 cells were measured for each condition. Statistical analyses were performed using Welch’s t-test with Bonferroni correction. *P<0.05, **P<0.01 and ***P<0.001.

Supplementary Figure S6. Resonance assignment of cell wall glucans in *C. neoformans* and RMSD comparison of β**-1,3-glucan polymorphs with model structures.**A) CP-based 2D ^13^C-^13^C correlation spectrum of rigid polysaccharides measured with 53 ms CORD mixing for KN99α and *fks1* mutants. (B) Through-bond carbon connectivity of mobile polysaccharides was resolved using 2D ^13^C DP refocused J-INADEQUATE spectrum. The absence of β-1,3-glucan signals in *fks1* mutants is shown by blue dotted boxes. (C) RMSD heatmap comparing ^13^C chemical shifts of polymorphic β-1,3-glucans with the model structures further supports the distinct conformational states of the two polymorphs. Stars indicate RMSD below 0.5 ppm. Note that type a closely resembles triple-helical β-1,3-glucan while type b exhibits a lower RMSD of 0.22 ppm relative to single-helical hydrated curdlan, indicating close structural correspondence with the single-helical conformation.

**Supplementary Figure S7. Genetic interaction scores in rich media without caspofungin treatment.** Rank-order waterfall plot showing genetic interaction score for mutants in each query background compared to wild type in YPAD. Highlighted genes are the same as for Fig 6B.

