## Supplementary material for "Therapeutic vulnerabilities of the drug-resistant fungal meningitis pathogen *Cryptococcus neoformans*": Figure S1

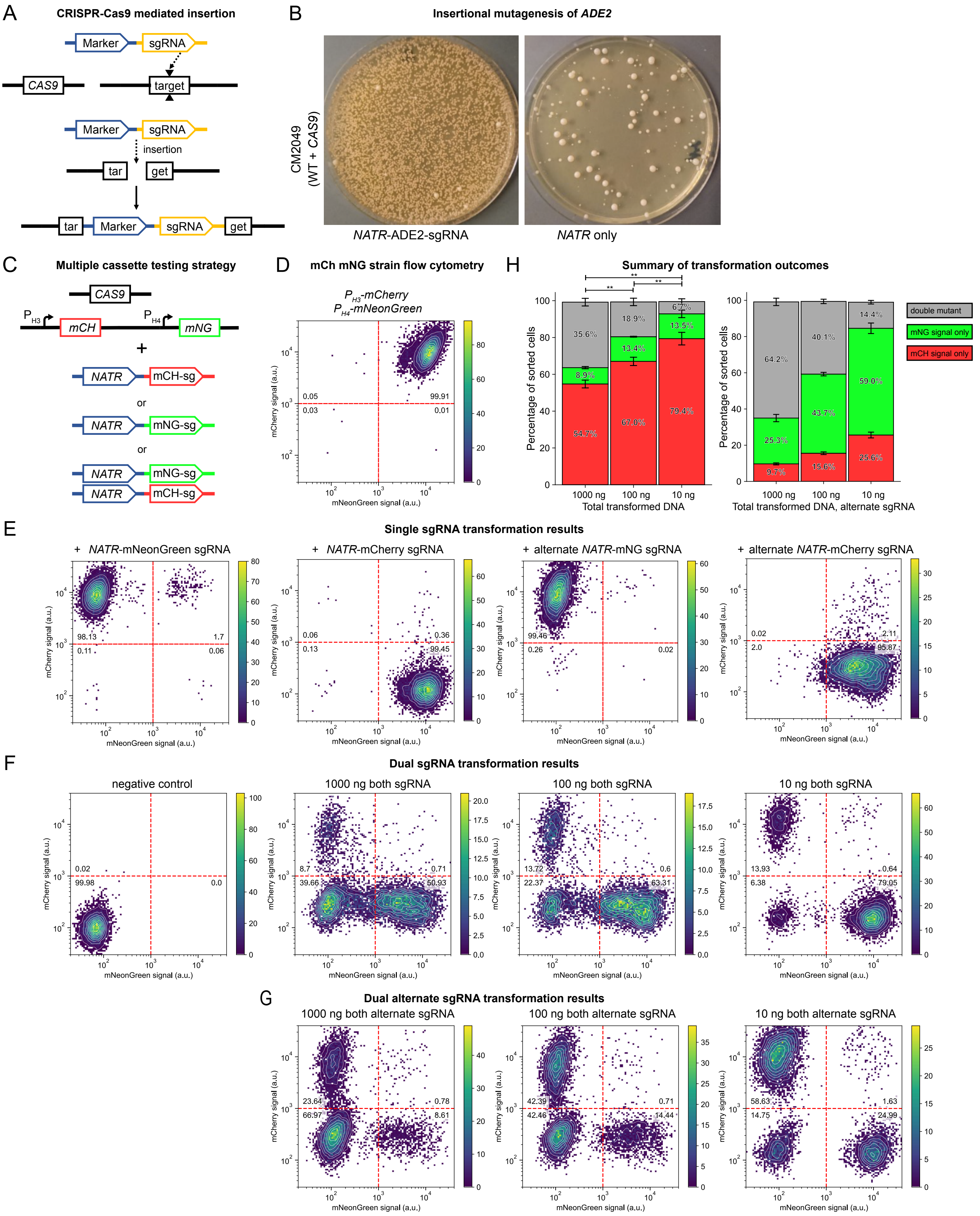

A
CRISPR-Cas9 mediated insertion
Marker
sgRNA
CAS9
target
insertion
tar
get
B
Insertional mutagenesis of ADE2
CM2049
(WT + CAS9)
NATR-ADE2-sgRNA
NATR only
C
Multiple cassette testing strategy
CAS9
P
H3
mCH
P
H4
mNG
+
NATR
mCH-sg
or
mNG-sg
D
mCh mNG strain flow cytometry
$P_{H3}$-mCherry
$P_{H4}$-mNeonGreen
0.05
99.91
0.03
0.01
mCherry signal (a.u.)
mNeonGreen signal (a.u.)
H
Summary of transformation outcomes
Percentage of sorted cells
35.6%
8.9%
54.7%
18.9%
13.4%
67.0%
6.7%
13.5%
79.4%
1000 ng
100 ng
10 ng
Total transformed DNA
64.2%
25.3%
9.7%
40.1%
43.7%
15.6%
14.4%
59.0%
25.6%
Total transformed DNA, alternate sgRNA
double mutant
mNG signal only
mCH signal only
E
Single sgRNA transformation results
+ NATR-mNeonGreen sgRNA
98.13
1.7
0.11
0.06
+ NATR-mCherry sgRNA
0.06
0.36
0.13
99.45
+ alternate NATR-mNG sgRNA
99.46
0.26
0.02
+ alternate NATR-mCherry sgRNA
0.02
2.11
2.0
95.87
F
Dual sgRNA transformation results
negative control
0.02
99.98
0.0
1000 ng both sgRNA
8.7
0.71
39.66
50.93
100 ng both sgRNA
13.72
0.6
22.37
63.31
10 ng both sgRNA
13.93
0.64
6.38
79.05
G
Dual alternate sgRNA transformation results
1000 ng both alternate sgRNA
23.64
0.78
66.97
8.61
100 ng both alternate sgRNA
42.39
0.71
42.46
14.44
10 ng both alternate sgRNA
58.63
1.63
14.75
24.99
