## Supplementary material for "Therapeutic vulnerabilities of the drug-resistant fungal meningitis pathogen *Cryptococcus neoformans*": Figure S2

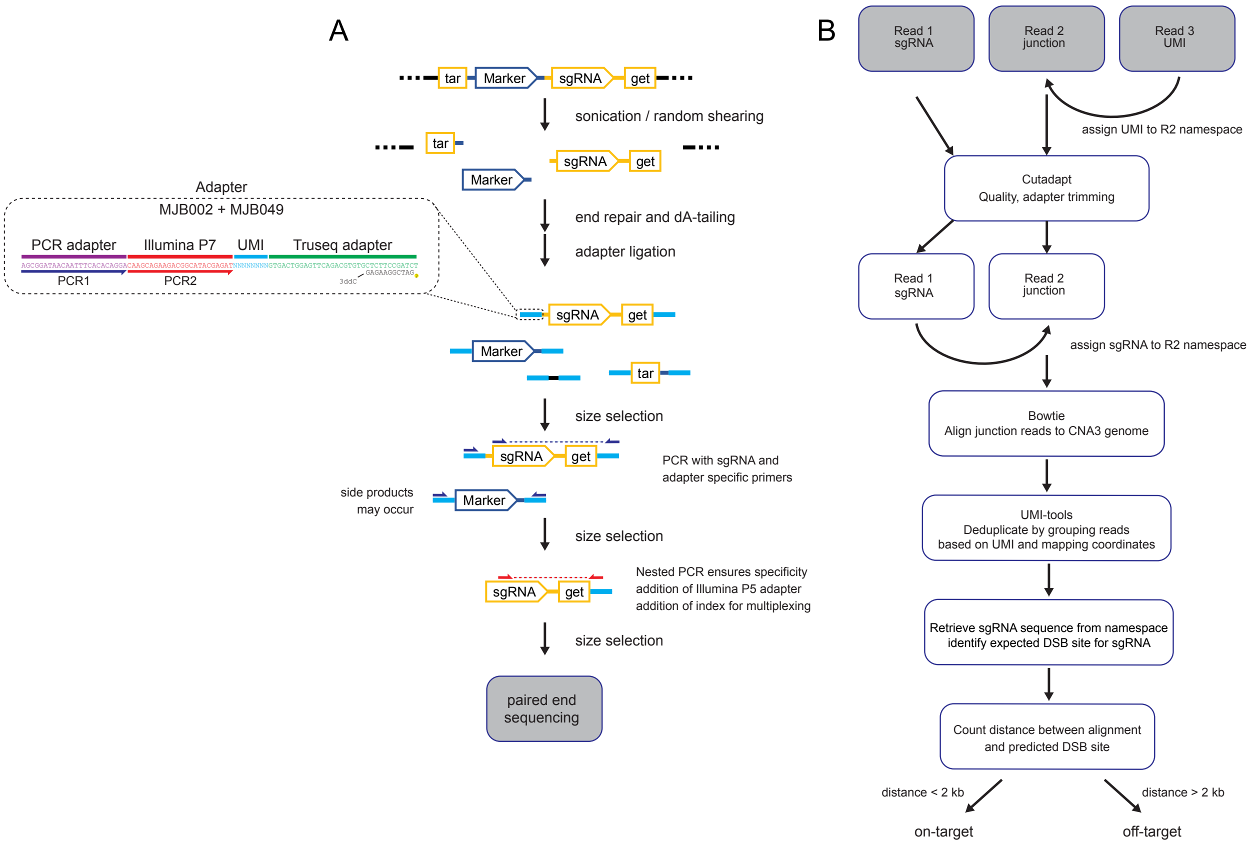
A
tar
Marker
sgRNA
get
sonication / random shearing
tar
sgRNA
get
Marker
Adapter
MJB002 + MJB049
PCR adapter
Illumina P7
UMI
Truseq adapter
AGCGGATAACAATTTCACACAGGACAAGCAGAAGACGGCATACGAGATNNNNNNNNGTGACTGGAGTTCAGACGTGTGCTCTTCCGATCT
GAGAAGGCTAG
3ddC
PCR1
PCR2
end repair and dA-tailing
adapter ligation
sgRNA
get
Marker
tar
size selection
sgRNA
get
PCR with sgRNA and adapter specific primers
side products may occur
Marker
size selection
sgRNA
get
Nested PCR ensures specificity
addition of Illumina P5 adapter
addition of index for multiplexing
size selection
paired end sequencing
B
Read 1 sgRNA
Read 2 junction
Read 3 UMI
assign UMI to R2 namespace
Cutadapt
Quality, adapter trimming
Read 1 sgRNA
Read 2 junction
assign sgRNA to R2 namespace
Bowtie
Align junction reads to CNA3 genome
UMI-tools
Deduplicate by grouping reads based on UMI and mapping coordinates
Retrieve sgRNA sequence from namespace identify expected DSB site for sgRNA
Count distance between alignment and predicted DSB site
distance < 2 kb
distance > 2 kb
on-target
off-target
