## Supplementary material for "Therapeutic vulnerabilities of the drug-resistant fungal meningitis pathogen *Cryptococcus neoformans*": Figure S3

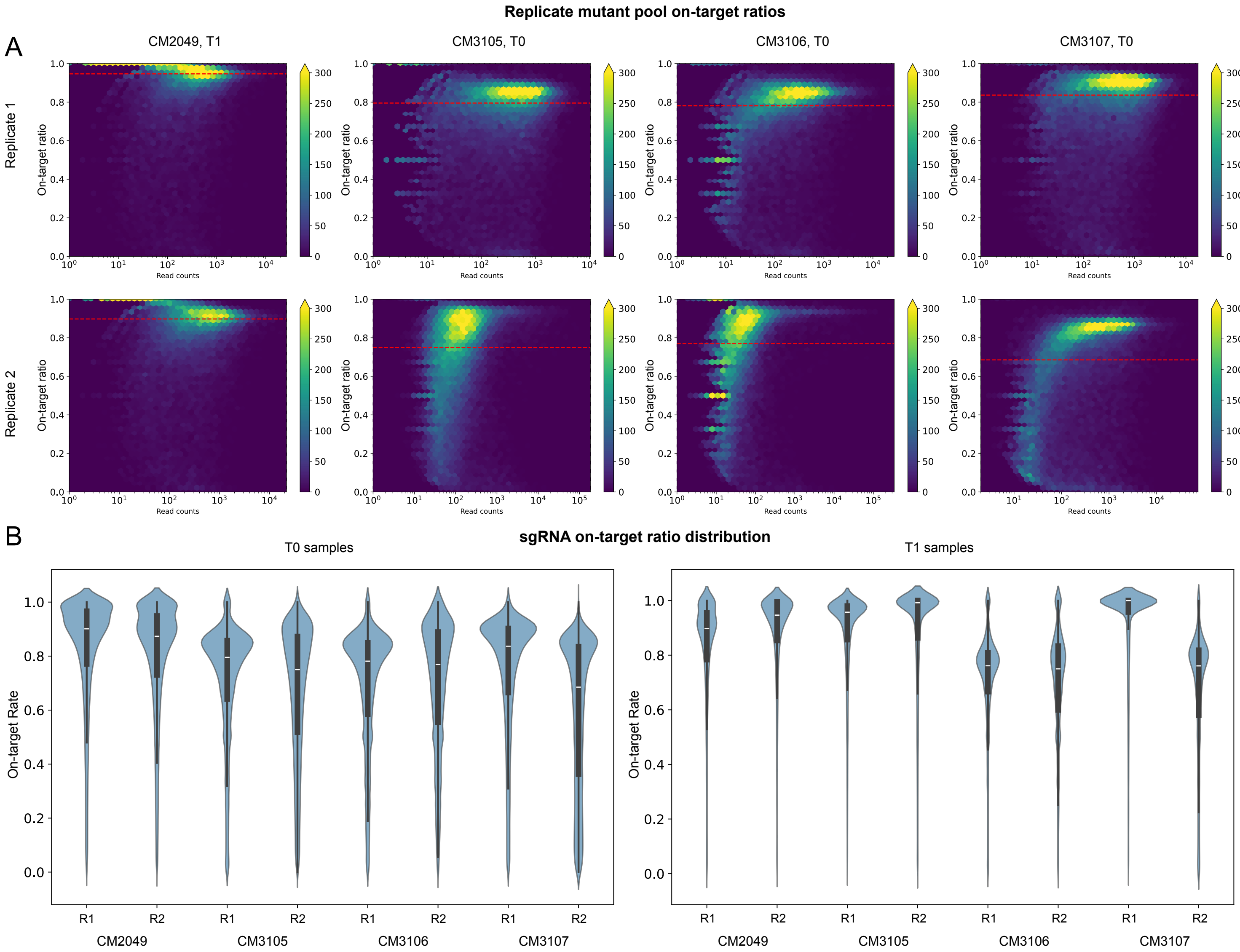
Replicate mutant pool on-target ratios
A
CM2049, T1
CM3105, T0
CM3106, T0
CM3107, T0
Replicate 1
Replicate 2
On-target ratio
Read counts
B
sgRNA on-target ratio distribution
T0 samples
T1 samples
On-target Rate
R1
R2
CM2049
CM3105
CM3106
CM3107
