## Supplementary figures and images for "Therapeutic vulnerabilities of the drug-resistant fungal meningitis pathogen *Cryptococcus neoformans*"

### Figure S4

#### Supplementary essential gene prediction testing

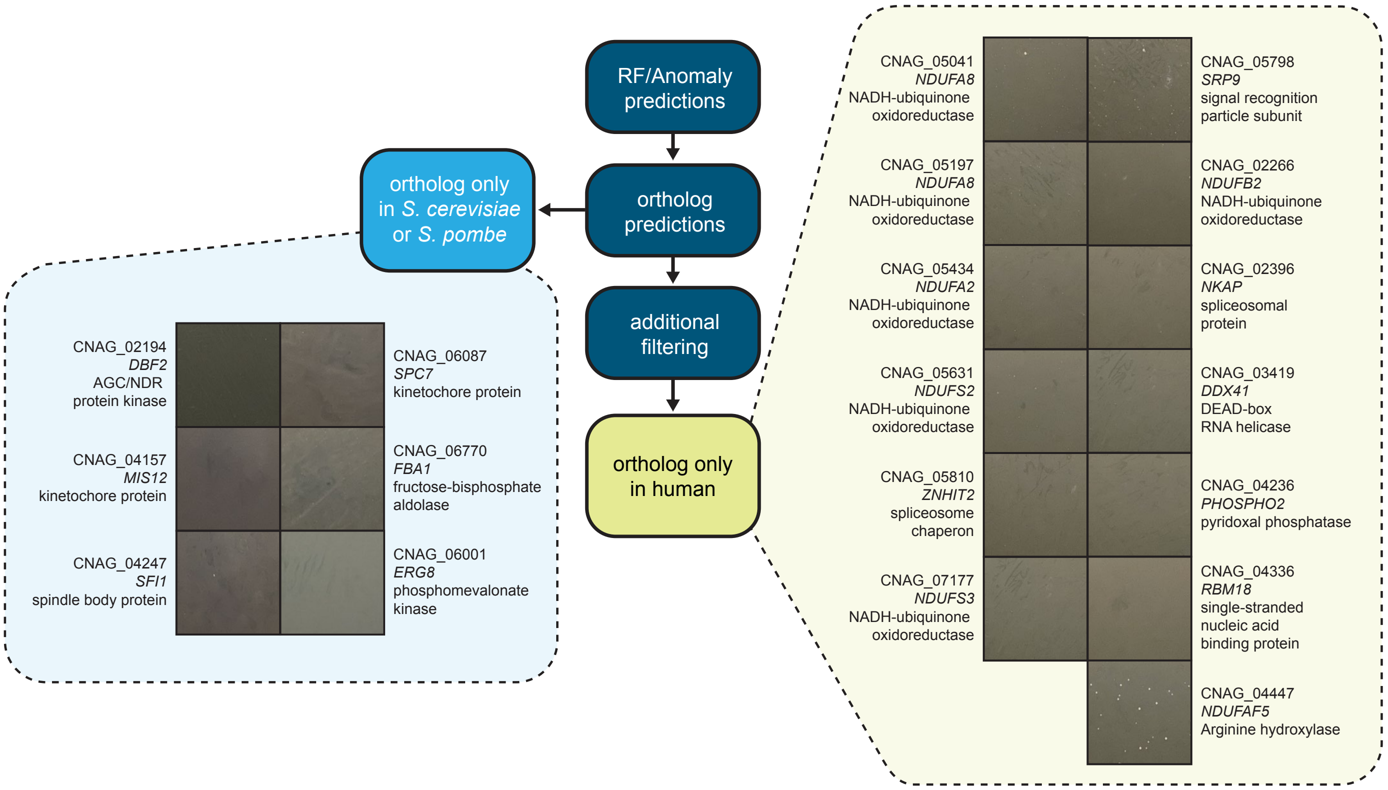

### Figure S7

#### Genetic interaction screening without caspofungin

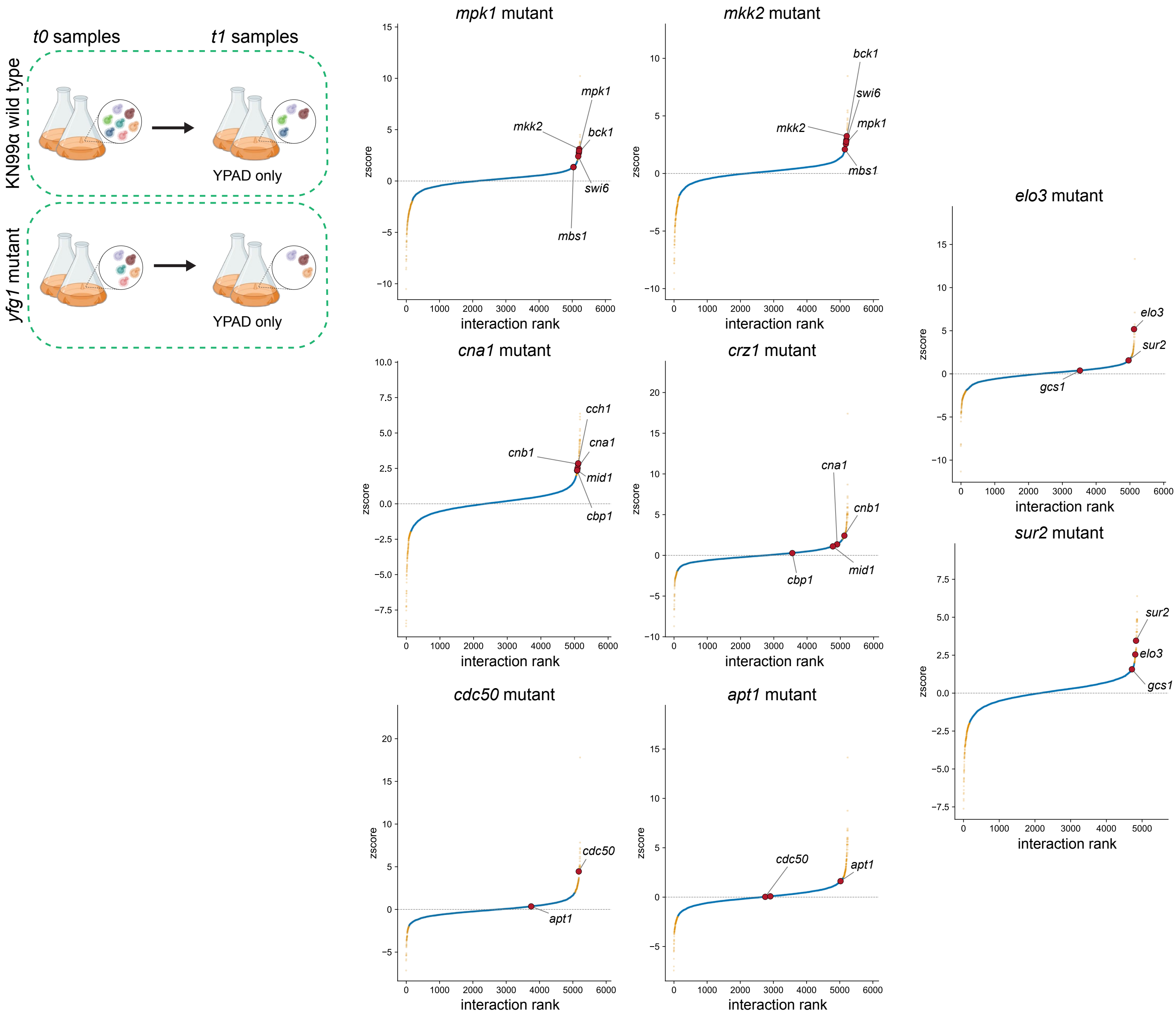
