## Supplementary material for "Therapeutic vulnerabilities of the drug-resistant fungal meningitis pathogen *Cryptococcus neoformans*": Figure S5

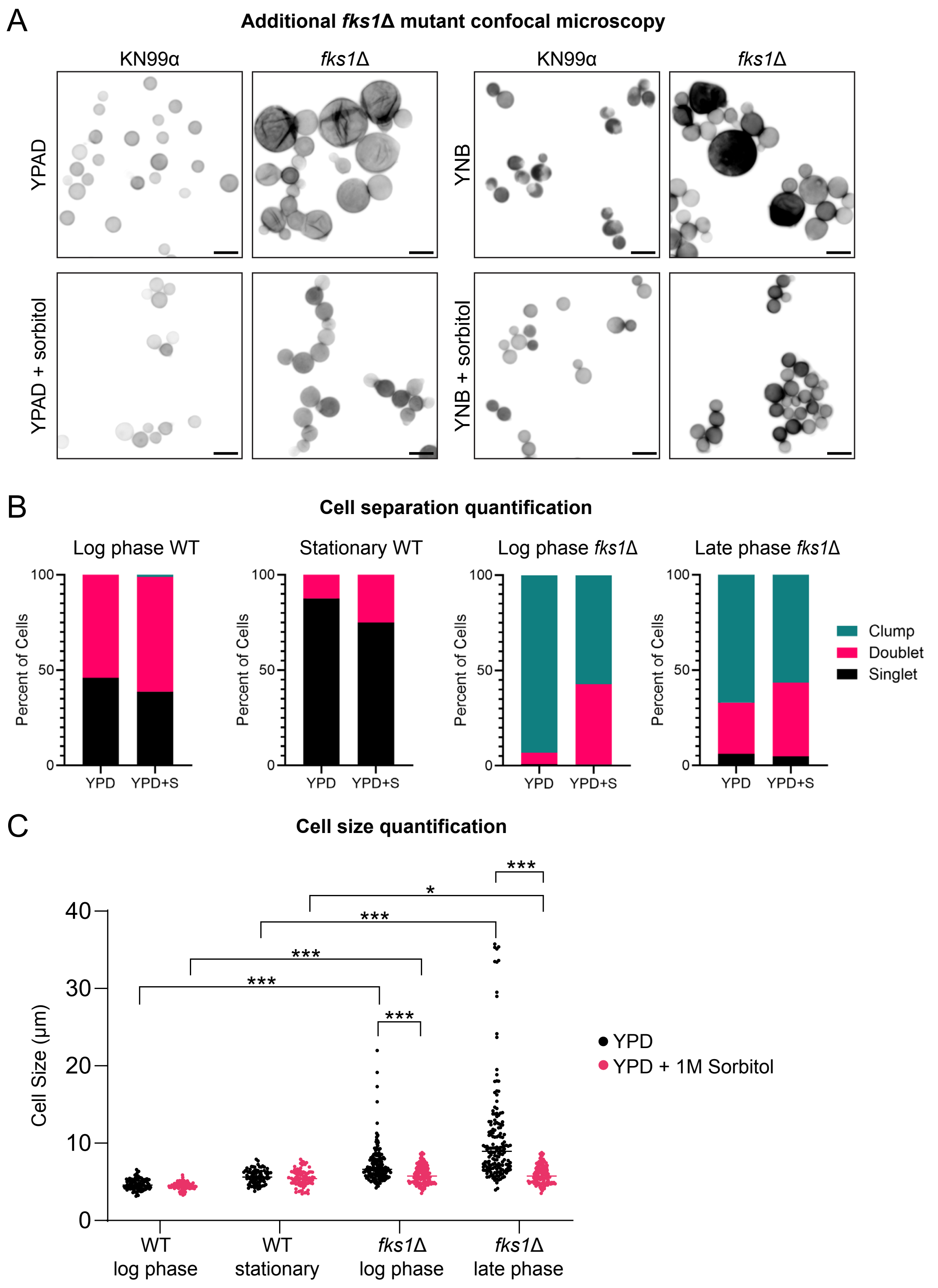
A
Additional fks1Δ mutant confocal microscopy
KN99α
fks1Δ
KN99α
fks1Δ
YPAD
YNB
YPAD + sorbitol
YNB + sorbitol
B
Cell separation quantification
Log phase WT
Stationary WT
Log phase fks1Δ
Late phase fks1Δ
Percent of Cells
100
50
0
YPD
YPD+S
Clump
Doublet
Singlet
C
Cell size quantification
Cell Size (µm)
40
30
20
10
0
***
*
YPD
YPD + 1M Sorbitol
WT
log phase
WT
stationary
fks1Δ
log phase
fks1Δ
late phase
