## Supplementary material for "Therapeutic vulnerabilities of the drug-resistant fungal meningitis pathogen *Cryptococcus neoformans*": Figure S6

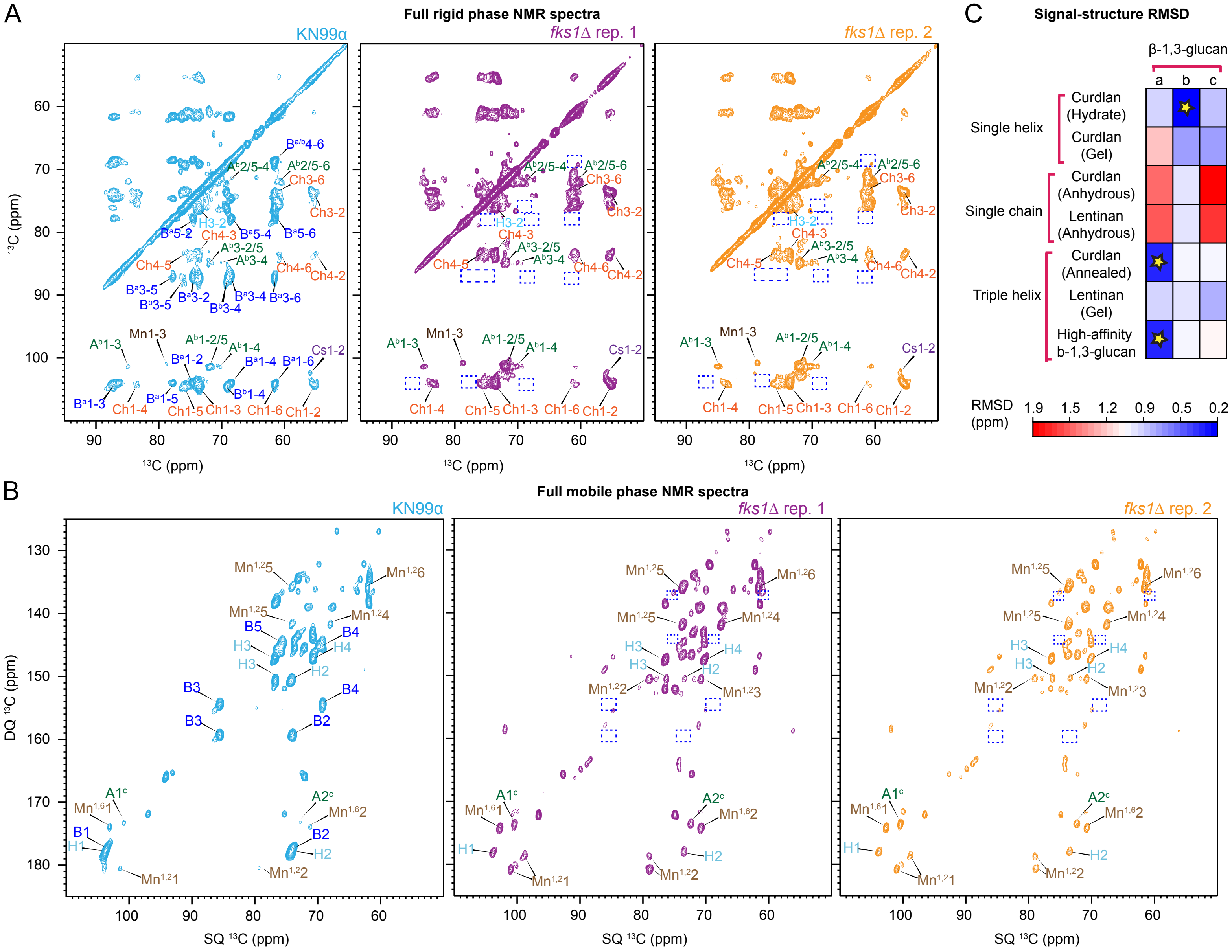
A
Full rigid phase NMR spectra
KN99α
fks1Δ rep. 1
fks1Δ rep. 2
13C (ppm)
13C (ppm)
Ba/b4-6
Ab2/5-4
Ab2/5-6
Ch3-6
Ch3-2
H3-2
Ba5-2
Ch4-3
Ba5-4
Ba5-6
Ab3-2/5
Ab3-4
Ch4-5
Ch4-6
Ch4-2
Ba3-5
Bb3-5
Ba3-2
Bb3-4
Ba3-4
Ba3-6
Ab1-3
Mn1-3
Ab1-2/5
Ab1-4
Ba1-2
Ba1-4
Ba1-6
Cs1-2
Bb1-4
Ba1-3
Ch1-4
Ba1-5
Ch1-5
Ch1-3
Ch1-6
Ch1-2
C
Signal-structure RMSD
β-1,3-glucan
a
b
c
Single helix
Curdlan (Hydrate)
Curdlan (Gel)
Single chain
Curdlan (Anhydrous)
Lentinan (Anhydrous)
Triple helix
Curdlan (Annealed)
Lentinan (Gel)
High-affinity b-1,3-glucan
RMSD (ppm)
1.9
1.5
1.2
0.9
0.5
0.2
B
Full mobile phase NMR spectra
KN99α
fks1Δ rep. 1
fks1Δ rep. 2
DQ 13C (ppm)
SQ 13C (ppm)
Mn1,25
Mn1,26
Mn1,24
B5
B4
H3
H4
H2
B3
B2
Mn1,22
Mn1,23
A1c
A2c
Mn1,61
Mn1,62
B1
H1
Mn1,21
Mn1,22
