## Supplementary Methods for "Therapeutic vulnerabilities of the drug-resistant fungal meningitis pathogen *Cryptococcus neoformans*"

1. **sgRNA double-joint PCR**
2. **sgRNA library cloning details**
3. **JuG-seq sequencing library preparation details**
4. **CRISPR-insert sequencing library PCR details**
5. **PCR of input DNA for CRISPR-insert**
1. **PCR conditions for double-joint PCR**

sgRNA PCR step 1:

Template is usually pBHM2329 1:100 dilution of miniprep (concentration not critical, use something low). Set up 2 reactions for each guide, 1 with sg_F + sgRNA_R1_R, 1 with sg_R + pcnu6_R1_F. Quantities are adjusted with excess for pipette error. Clean both products on the same column and elute in 40 µL sterile water.

| 10x Extaq Buffer | 5 µL |
| --- | --- |
| 2.5mM Extaq Kit dNTPs | 4 µL |
| 10 uM F' | 2 µL |
| 10 uM R' | 2 µL |
| ddH2O | 37 µL |
| Template | 1 µL |
| Extaq pol. | 0.2 µL |

| 98°C | 2:00 |
| --- | --- |
| 98°C | 0:15 |
| 52°C | 0:30 |
| 72°C | 0:30 |
| Go to 2 | 34x |
| 72°C | 5:00 |
| 4°C | hold |

sgRNA PCR step 2:

Using the round 1 product as template, set up a PCR as follows:

| 10x Extaq buffer | 5 µL |
| --- | --- |
| 2.5mM Extaq Kit dNTPs | 4 µL |
| dH2O | 31 µL |
| Template | 10 µL |
| Extaq pol | 0.25 µL |

| 94°C | 3:00 |
| --- | --- |
| 94°C | 0:30 |
| 55°C | 10:00 |
| 72°C | 5:00 |
| Go to 2 | 9x |
| 72°C | 10:00 |
| 4°C | hold |

sgRNA PCR step 3:

Using nested primers (M13/F and M13/R usually) set up PCR with the same conditions as round 1, using the round 2 product as template. Clean products on column per manufacturer’s instructions, elute products in sterile water.

Marker PCR:

Amplification of selectable markers are set up as follows, using 1-10 ng of the appropriate plasmid template. Clean products on column per manufacturer’s instructions, elute products in sterile water.

| 10x Extaq Buffer | 5 µL |
| --- | --- |
| 2.5mM Extaq Kit dNTPs | 4 µL |
| 10 uM F' | 2 µL |
| 10 uM R' | 2 µL |
| ddH2O | 34.5 µL |
| DMSO | 2.5 µL |
| Template | 1 µL |
| Extaq pol. | 0.2 µL |

| 94°C | 3:00 |
| --- | --- |
| 94°C | 0:30 |
| 50°C | 0:30 |
| 72°C | 3:00 |
| Go to 2 | 34x |
| 72°C | 5:00 |
| 4°C | hold |

2. **sgRNA library cloning details**

The sgRNA library was ordered as an oligo pool, diluted to 10 ng/µL, and amplified using P01 and P02 as follows.

| 2x NEBNext Q5 MM | 50 µL |
| --- | --- |
| 100 µM F' | 1 µL |
| 100 µM R' | 1 µL |
| 10 ng/µL oligo pool | 2 µL |
| ddH2O | 46 µL |

| 98°C | 3:00 |
| --- | --- |
| 98°C | 0:15 |
| 55°C | 0:15 |
| 72°C | 0:20 |
| Go to 2 | 15x |
| 72°C | 2:00 |
| 4°C | hold |

DNA is then cleaned on PCR cleanup columns per manufacturer’s instructions and eluted in 10 µL sterile water. The PCR product is then cloned in a gibson/assembly reaction as follows.

| Digested, gel purified vector | 500 ng |
| --- | --- |
| PCR amplified library | 25 ng |
| Sterile water | to 30 µL |
| NEB 2x Hifi Assembly MM | 30 µL |

Incubate for 1 hour at 50°C. Next, samples are cleaned using Biorad P-30 buffer exchange columns. Columns are equilibrated 5x with 500 µL sterile water with 1 min spins at 1000g. The total product from the assembly reaction is loaded onto the column and spun for 4 min at 1000g. Volume is then adjusted to 100 µL in water, and 2 µL GlycoBlue, 10 µL 3M pH5.2 NaOAc, and 300 µL 100% ethanol are added to the sample. The sample is then incubated at -20°C overnight before being spun at 18,000 g for 30 minutes at 4°C. Pellet is washed twice in cold ethanol, allowed to dry, then resuspended in 10 µL sterile water for electroporation.

3. **JuG-seq sequencing library preparation details**

Shearing, end-repair, adapter ligation, are all conducted per manufacturer’s instructions and are described in Methods. To anneal the adapter, 12 µL each of 200 µM P04 and P05 are combined, mixed by vortexing, then heated in a thermocycler to 95°C, held for 5 minutes, and cooled to 12°C with ramp speed set to 0.1°C/s. Annealed adapter is then diluted to 15 µM by adding 136 µL of sterile water and this product is then used in 20 µL aliquots.

After adapter ligation and size selection using Ampure XP beads, samples are PCR amplified in two rounds. Conditions for both rounds are identical and are as follows.

| 2x NEBNext Q5 MM | 25 µL |
| --- | --- |
| 10 µM F' | 5 µL |
| 10 µM R' | 5 µL |
| Ligated or Round 1 product | 15 µL |

| 98°C | 0:30 |
| --- | --- |
| 98°C | 0:10 |
| 65°C | 1:30 |
| Go to 2 | 15x |
| 65°C | 5:00 |
| 4°C | hold |

4. **CRISPR-insert sequencing library PCR details**

For amplification of the sgRNA and transformant UMI, PCR conditions are as follows.

| 2x NEBNext Q5 MM | 50 µL |
| --- | --- |
| 100 µM F' | 1 µL |
| 100 µM R' | 1 µL |
| Sample genomic DNA | 500 ng |
| ddH2O | To 100 µL |

| 98°C | 3:00 |
| --- | --- |
| 98°C | 0:30 |
| 56°C | 0:15 |
| 72°C | 0:30 |
| Go to 2 | 14x |
| 72°C | 3:00 |
| 4°C | hold |

5. **PCR of input DNA for CRISPR-insert**

PCR conditions as follows:

| 5x Q5 buffer | 20 µL |
| --- | --- |
| 2.5mM dNTPs | 8 µL |
| 10 uM F' (M13/F) | 5 µL |
| 10 uM R' (M13/R) | 5 µL |
| ddH2O | 42 µL |
| Q5 enhancer | 20 µL |
| Template (10 ng/µL) | 2 µL |
| Q5 polymerase | 1 µL |

| 98°C | 2:00 |
| --- | --- |
| 98°C | 0:20 |
| 52°C | 0:30 |
| 72°C | 1:20 |
| Go to 2 | 25x |
| 72°C | 5:00 |
| 4°C | hold |
